# epiGBS2: an improved protocol and automated snakemake workflow for highly multiplexed reduced representation bisulfite sequencing

**DOI:** 10.1101/2020.06.23.137091

**Authors:** Fleur Gawehns, Maarten Postuma, Thomas P. van Gurp, Niels C. A. M. Wagemaker, Samar Fatma, Morgane Van Antro, Christa Mateman, Slavica Milanovic-Ivanovic, Kees van Oers, Ivo Grosse, Philippine Vergeer, Koen J. F. Verhoeven

## Abstract

epiGBS is an existing reduced representation bisulfite sequencing method to determine cytosine methylation and genetic polymorphisms *de novo*. Here, we present epiGBS2, an improved epiGBS laboratory protocol and user-friendly bioinformatics pipeline for a wide range of species with or without reference genome. epiGBS2 decreases costs and time investment and increases user-friendliness and reproducibility. The library protocol was adjusted to allow for a flexible choice of restriction enzymes and a double digest. Instead of fully methylated adapters, semi-methylated adapters are now used. The bioinformatics pipeline was improved in speed and integrated in the snakemake workflow management system, which now makes the pipeline easy to execute, modular, and parameter settings flexible. We also provide a detailed description of the laboratory protocol, an extensive manual of the bioinformatics pipeline, which is publicly accessible on github (https://github.com/nioo-knaw/epiGBS2) and zenodo (https://doi.org/10.5281/zenodo.3819996), and example output.

## Introduction

Cytosine methylation at carbon position 5 (also termed 5-meC) is a chemical epigenetic modification of DNA. This modification can influence gene activity and expression and has the potential to affect transcription regulation. Genome-wide 5-meC discovery is routinely performed by using methods based on bisulfite treatment followed by high throughput sequencing (BS-Seq)^1^. Whole genome BS-Seq (WGBS)^2^ is the golden standard if the financial resources and a reference genome are available, which is still not the case for the majority of organisms. While the popularity of bisulfite sequencing is growing (supplemental table), data are mainly generated for model species like human, mouse and *A. thaliana* representing 42%, 35% and 6% of all BS-Seq data sets in the SRA (by September 2019), respectively.

A less comprehensive but cheaper and versatile alternative to WGBS is BS-Seq in reduced representations of the genome by using restriction enzyme fragmentation during the library preparation (e.g. RRBS^3^, epiGBS^4^ and BsRADseq^5^ or epiRADseq^6^). Several easy-to-use bioinformatics tools and workflows are developed to analyze BS-seq data, such as BS-Seeker2^7^, Bismark^8^ and BAT^9^. However, the interest in understanding the significance of epigenetics in ecology and evolution increases and requires methods, which can handle genomes of evolutionarily divergent species as well as for DNA methylation analysis in species for which a reference genome is lacking. Such methods have to deal with the complex genomes of non-model organisms, high sample numbers and accommodate a simultaneous comparison of genetic and epigenetic data, for instance to examine how much of the overall epigenetic variation between samples can be predicted from pairwise genetic relatedness^10^.

In a previous publication, we presented epiGBS as a reduced-representation DNA methylation analysis tool that combines those features^4^. EpiGBS calls both cytosine-level quantitative DNA methylation scores and SNPs from the same bisulfite-converted samples, while reconstructing the *de novo* consensus sequence of the targeted genomic loci. This means that the method can be applied also when no reference genome is available for the species under study^4^. Here, we present an update of the epiGBS laboratory and computational analysis protocols. Compared to the original epiGBS method, the updated protocols in epiGBS2, as presented here, decrease costs and time investment and increase user-friendliness and reproducibility, also allowing for an effective use of epiGBS in a wider range of species, including vertebrates such as birds^11^.

## Modifications to the lab protocol

In order to reduce sequencing bias and costs, several major improvements were made to the epiGBS laboratory protocol (supplemental materials), which were described recently by Boquete *et al.*^12^. We briefly list the key improvements here; In addition, we present a detailed description of the adapter design, which allows free choice of the restriction enzyme pair.

### Identification of PCR duplicates

During the preparation of sequencing libraries, PCR clones can be produced. Removing these PCR duplicates computationally removes overrepresented fragments caused by biased duplication, which allows for more accurate interpretation of results. Using common whole-genome sequencing laboratory protocols, sequence identity is a basis for distinguishing PCR duplicates. However, in reduced representation approaches that use amplification of restriction enzyme-associated DNA, fragments of identical sequence are produced by design from starting DNA; sequence identity is therefore not a basis for distinguishing PCR duplicates. To differentiate PCR duplicates from epiGBS sequencing reads that originate from different DNA molecules, a random three letter oligonucleotide was placed in the adapter sequence as described in van Moorsel, *et al*^13^ (Fig. 1). This Unique Molecular Identifiers (UMI) or so-called “Wobble” sequence is identical for PCR clones but different for true replicates. This feature is used in the epiGBS computational workflow to specifically remove PCR clones.

**Figure 1:**
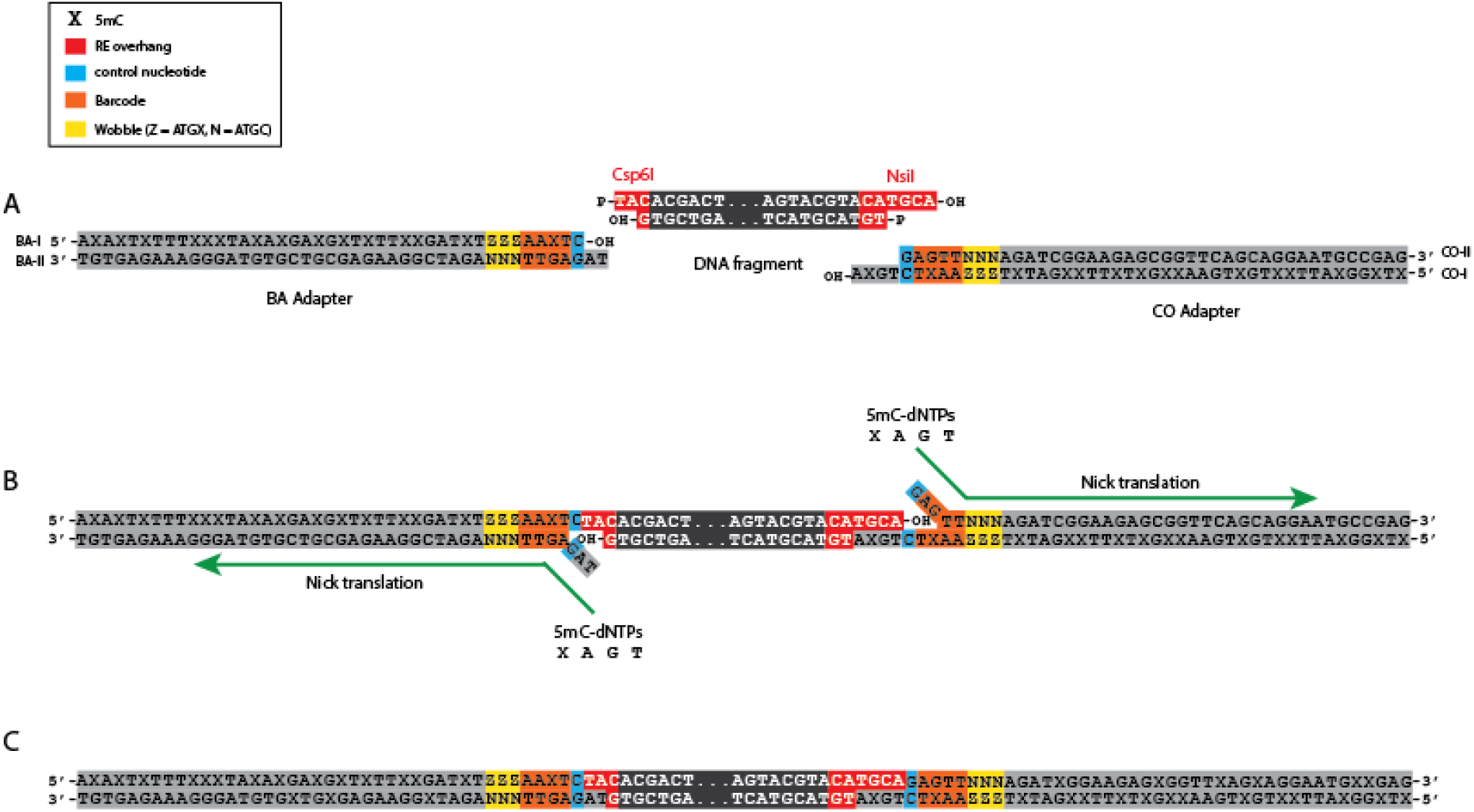
epiGBS2 uses hemi-methylated adapters. (A) The 5’-3’ strand of the BA adapter and the 3’-5’ strand of the CO adapters are made containing methylated cytosines (X); the opposite strands are unmethylated. (B) All strands are de-phosphorylated, so only adapter 3’ ends and DNA fragment 5’ ends ligate. During Nick-translation the “broken” strands are replaced with 5mC-dNTPs, which results in fully methylated adapters (C). Yellow = random three nucleotides (UMI/”Wobble”), orange = barcode, blue = control nucleotide, red = restriction enzyme overhang

### Use of a control nucleotide and an universal two restriction enzyme digest

The epiGBS protocol in van Gurp, *et al.*^4^ used a single restriction enzyme (RE) digest with either *PstI* or *Csp6I*. Both RE recognition sites contained a cytosine that either originated from the adapters BA and CO (methylated C, unconverted) or from the DNA fragment (unmethylated C, converted) in the sequencing reads. Fragments with unconverted recognition sites on barcode adapter BA were arbitrarily defined as Watson; fragments with converted recognition sites on barcode adapter CO as Crick. We now added the possibility to perform a double RE digest, e.g. with a rare and a frequent cutting RE. To allow the use of REs without a C in their recognition site, a “control nucleotide” (CN) was introduced in the adapter (see van Moorsel, *et al*^13^ for a further description). This CN is an un-methylated cytosine, which is placed after the barcode followed by the sequence of the RE overhang (Fig. 1) and used for Watson/Crick annotation of the reads. Read pairs with T at the CN position of the R1/ adapter BA read and C at the CN position of the R2/ adapter CO read are defined as Watson; read pairs with C at the CN position of the R1/ adapter BA read and T at the CN position of R2/ adapter CO read as Crick. This design facilitates the usage of various RE combinations and makes the epiGBS protocol more universally applicable. While the epiGBS protocol was originally optimized for plants, the freedom to also use other enzymes, such as for example *MspI*, makes epiGBS2 now also very effective for studies on other organisms, such as vertebrates.

### Use of hemi-methylated adapters

By highly multiplexing samples, and using a GBS-based protocol with custom barcoded adapters, the costs of epiGBS are low in comparison to common RRBS approaches. However, the use of fully methylated adapters is relatively expensive, depending on the vendor. To further reduce costs, the library preparation protocol was adjusted in such a way that hemi-methylated adapter pairs are used instead of fully methylated adapters. In epiGBS2 the cytosines of the oligonucleotides Adapter BA-I and Adapter CO-I are 5-C methylated (Fig. 1 and see laboratorial protocol in the supplemental materials). The oligonucleotides of the opposite strands (Adapter BA-II and Adapter CO-II) contain un-methylated cytosines only and are 5’-de-phosphorylated. After annealing the respective BA-I and BA-II and CO-I and CO-II adapter oligonucleotides and ligating them with the enzyme digested DNA fragment, only adapter 3’ ends and fragment 5’ ends ligate. A nick remains between adapter 5’ ends and fragment 3’ ends. The nick is repaired by using dNTPs that contain 5-meC’s and that directly translates all 5’-3’ nucleotides starting from the nick. This results in fully methylated adapters that are ligated to the digested DNA fragment and a complementary short UMI/ “Wobble” sequence.

## Improvements in the computational analysis protocol

The epiGBS analysis scripts of van Gurp et al. (2016) were updated with the aims to improve performance, user-friendliness and reproducibility of analysis.

### Embedding into Snakemake workflow and conda

Workflow management systems (WMS), such as nextflow^14^ or snakemake^15^, are a way of describing analytical pipelines and computational tools. These systems have a common aim: to make computational methods reproducible, portable, maintainable and shareable. WMS make sure to monitor the progress of e.g. bash or python scripts and exit gracefully if any step fails^16^. WMS also integrate with package managers like conda and Docker, which install software dependencies into the working directory, without requiring any admin/root privileges. In combination, WMS and package managers make the installation and execution of analysis pipelines accessible for biologists who have a basic knowledge in bioinformatics. The computational epiGBS workflow consists of seven python scripts, which we here embedded in a snakemake workflow (Fig. 2). Each script is called in a specific snakemake rule resulting in modularity of the bioinformatics analysis. Hence, the pipeline can run all steps at once or could be executed in specific parts.

**Figure 2:**
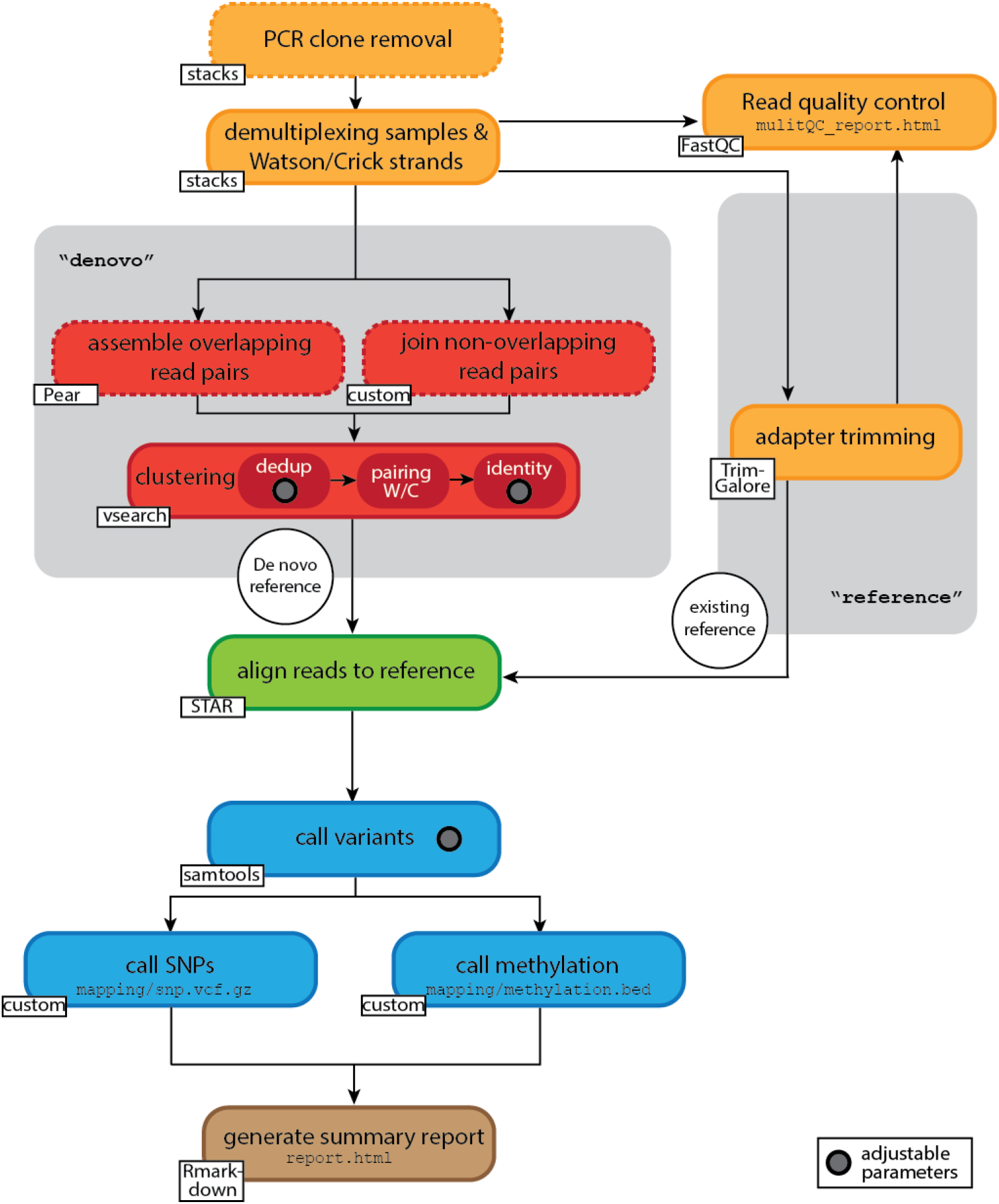
Overview of the epiGBS2 modules. The main snakemake modules are visualized. Boxes with solid lines represent steps with modular output and can be executed independently; steps in boxes with dashed lines are not modulated. Orange: Read preprocessing. PCR clones are removed from all input reads, samples are demultiplexed with stacks and annotated as either Watson or Crick reads. In the reference branch reads are additionally trimmed with Trim-Galore/cut-adapt^26^. The processed reads are analysed in a read quality control using FastQC^28^ and summarized with MultiQC^25^. Red: In the de novo branch reads are either assembled with Pear^29^ or joined with a custom script. These sequences are deduplicated, Watson and Crick reads paired and clustered based on identity. The minimum cluster size during deduplication and the identity percentage are introduced as variable parameters and can be set in the config file. Green: Reads are aligned to the reference (either de novo clusters or pre-existing reference) with STAR^22^. Blue: Variants are called with samtools mpileup^24^ and processed by custom scripts to identify SNPs and methylations sites. In samtool mpileup the maximum read depth can be varied. Brown: A summary report is generated containing important quality measurements for the analysis with Rmarkdown.

To improve the installation of the pipeline, we created conda^17^ environments in such a way that the workflow is portable and independent from the used Linux system. Each snakemake rule calls specific conda environment files and automatically installs the required dependencies.

### Demultiplexing

epiGBS2 takes the raw sequencing reads and a barcode file as input. First, the PCR clones, which are identified by identical sequence and UMI/ “Wobble” sequence, are removed. Then reads are demultiplexed without allowing any mismatches according to their barcode and control nucleotide sequence. Only reads with confirmed presence of the expected restriction enzyme overhang are retained. Read headers are labelled with their sample identification code and with either “Watson'' or “Crick”. While the original epiGBS pipeline used custom scripts, these processes are now executed by the *filter_clone* and *process_radtags* commands of the Stacks 2^18–20^software. Consequently, the speed of the demultiplexing was increased by a factor of approximately six.

### Mapping

Mapping was previously performed with bwa-meth^21^ but is now implemented with the fast alignment program STAR^22^ (version 2.5.3). For this purpose, the read files and the reference sequences are converted with custom scripts into a Watson (A, T, G) - or Crick (A, C, T)-dependent three letter alphabet. The STAR parameters were adjusted to meet the requirements for aligning BS treated reads (Tabel 1) with reduced nucleotide diversity:

1. In epiGBS only reads that map uniquely are kept.
2. The alignment type was set to EndToEnd and soft clipping is prevented.
3. Gap penalty was increased in general, but decreased for AT/AC, GT/AT and non-canonical junctions.
4. The ratio of mismatches to mapped length was adjusted.

**Tabel 1:**
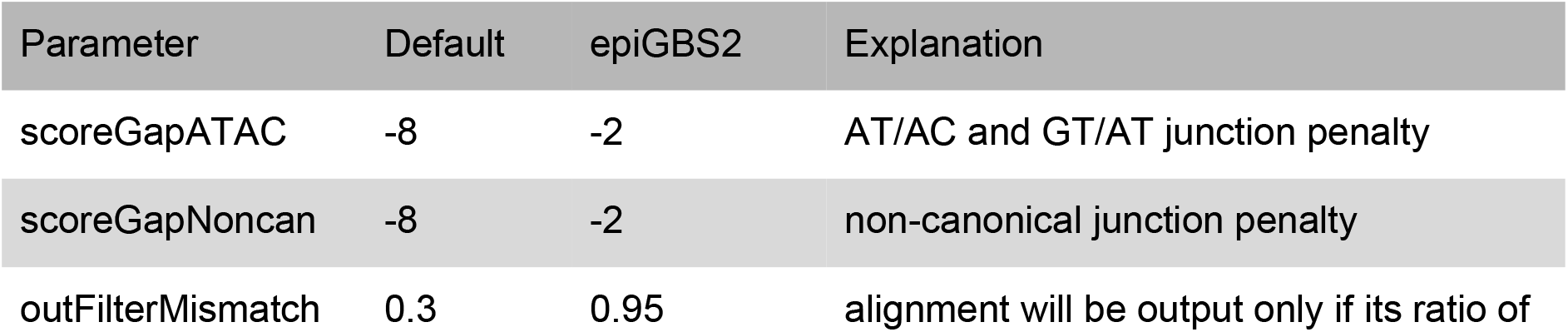
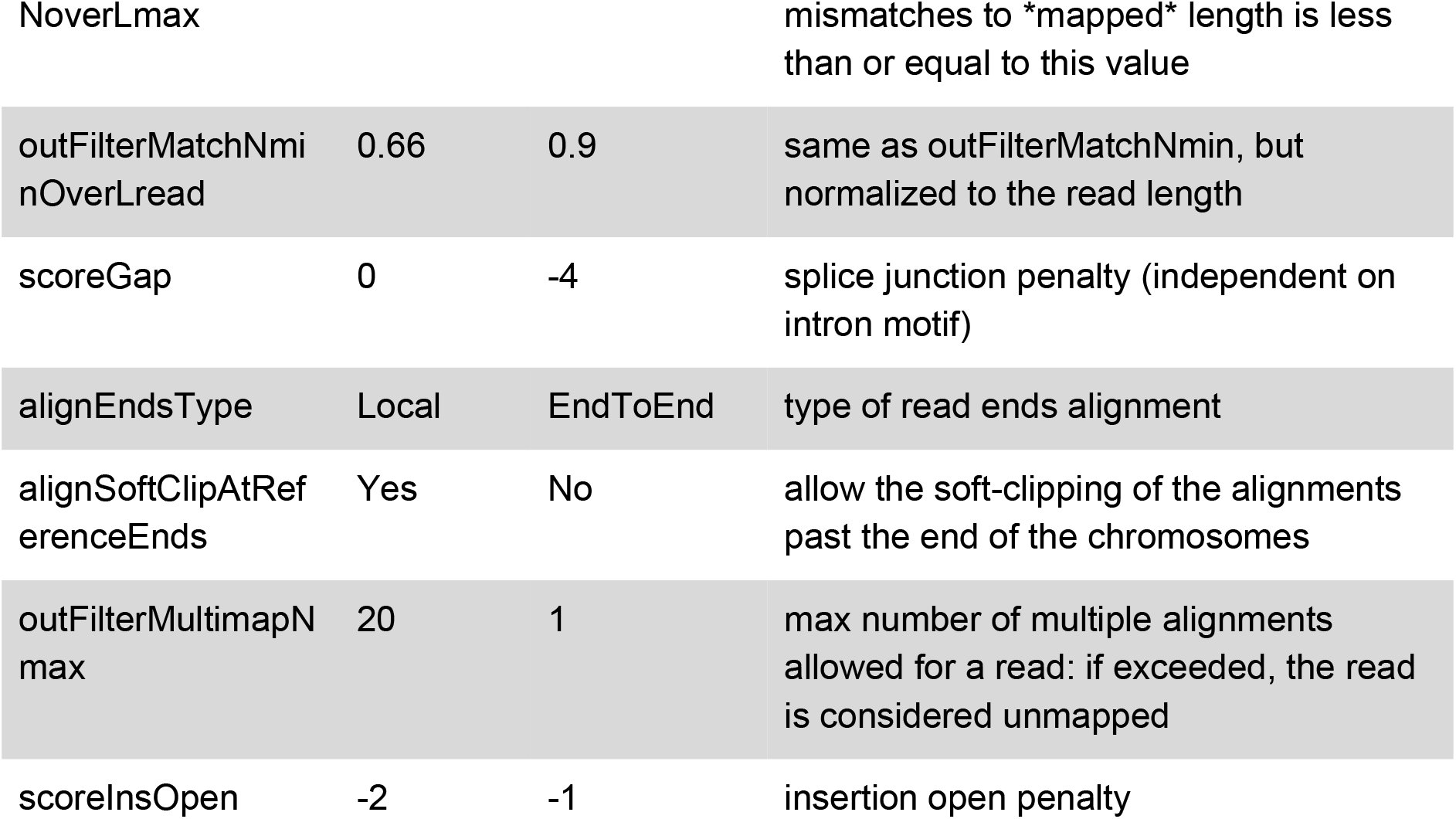
Parameters of the alignment program STAR. To enable read mapping with STAR, parameters were changed from default to epiGBS2-specific. This table shows the values of changed parameters and a short explanation.

### SNP / Methylation calling

Previously, SNP and methylation calling were implemented by polymorphism discovery with Freebayes^23^, followed by differentiating between genetic and epigenetic variation by comparing the nucleotide sequence of Watson and Crick strands^4^.

EpiGBS2 now uses custom scripts to call methylation variants and SNPs. First, a "pileup" textual format is generated from the Watson and Crick alignment using samtools mpileup^24^ (version 1.3.1). Then Watson and Crick variant information is merged into a single file, from which SNP and methylation sites are called by custom scripts. The custom SNP calling algorithm compares the reference nucleotide with the calls in the Watson reads and the Crick reads. A possible methylation site is identified when the reference nucleotide is a guanine (G) or a cytosine (C) and the called nucleotide a G and an adenosine (A) or a thymine (T) and a C in the Watson and Crick strand, respectively.

### Creation of a summary output file

The workflow produces two report files: A report summarizing the read quality of the processed reads using MultiQC^25^ (version 1.8) and another report (example file in the supplemental material) summarizing the statistics of the read processing like clone removal, demultiplexing and trimming, *de novo* reference re-construction, mapping and variant calling. The latter was implemented using custom R (version 3.6.1) and python code in Rmarkdown and rendered with knitr (version 1.22). Plots include visualization of the cloned read counts, distribution of the read counts per sample, SNP depth per sample, called samples per methylation site, methylation site depth per sample, number of methylation sites per context (CG, CHG, CHH) and methylation ratio counts per methylation context. All methylation-specific plots are created based on the first 100.000 positions of the methylation output file (methylation.bed) to save memory usage and generation time; thus, these plots are meant to provide a quick visual check of the pipeline results but are not intended to reflect final results. Summary statistics are extracted from different log files and give an overview of the assembly efficiency of the paired reads, the number of created *de novo* reference sequences (clusters) and mapping efficiency.

### Implementing a reference genome branch

epiGBS2 runs in two modes: either with a pre-existing reference genome or in a *de novo* mode. In *de novo* mode, the reference of the fragments under study is reconstructed from the epiGBS reads^4^. A reference mode was added to the workflow, which did not exist previously but which facilitates the use of epiGBS when a reference genome is available for the study species.

Adapter trimming was introduced as a major adjustment to enable implementation of the reference mode. This was needed because the majority of the de-duplicated and demultiplexed paired-ends reads are longer than the DNA fragment length, and hence carry adapter sequences at the 3’-end. In *de novo* mode, those adapter sequences are removed by merging the read pairs. In the reference mode, however, adapter trimming is executed using Trim Galore! (version 0.5.0) and cutadapt^26^, which by default recognizes the commonly used Illumina adapters. This is followed by additional trimming of the first 10 bp at the 3’-end of the reads to remove the custom-made parts of the adapters, including the three random nucleotides, the barcodes and the control nucleotide. Trimming is followed by mapping and variant calling as performed for the *de novo* mode.

### Flexible parameter setting for de novo reference sequence reconstruction and for SNP calling

We added the possibility to vary a number of parameters directly from the config file, for increased flexibility during analysis. These modifications are implemented in two of the epiGBS pipeline modules (Fig. 2). 1) *De novo* reference construction. The reconstruction of the *de novo* sequence consists of three clustering steps: a) deduplication of three-letter encoded Watson and Crick-reads; b) pairing Watson and Crick-reads; and c) clustering of reconstructed reference clusters by identity. The performance of a) can now be customized by setting a minimum and/or maximum cluster depth, and c) can be customized by setting the identity % for clustering. 2) Variant calling. The first step in the calling of SNPs and methylation variants creates a variant file in pileup format that summarizes the number of reads covering a specific site in the reference, the read bases and qualities. During creation of this file, samtools decreases the number of reads at highly covered positions in order to avoid exceeding memory constraints of the system. This can now be manipulated by setting the maximum-depth parameter. Other parameter values that can be customized in this step are minimal mapping quality and a minimal base quality.

### Miscellaneous improvements

- As support of Python 2 stopped on January 1, 2020 (https://www.python.org/doc/sunset-python-2/) all python scripts were transferred to python 3.
- Before mapping reads to the reference, all sequences are converted from a four-letter alphabet (ACTG) to a three-letter alphabet (ATG for Watson or CGT for Crick). In the original code, the last reference cluster in the reference sequence file was missed and not taken into account during mapping. This error was fixed: in epiGBS2 all reference clusters are considered.
- The header of the file *merged.tsv* was fixed and changed from: Chromosome (CHROM), position (POS), ID, reference, alternative (ALT), quality (QUAL), FILTER, INFO, FORMAT, sample names to: Chromosome (CHROM), position (POS), reference (REF), alternative Watson allele (ALT_WATSON), alternative Crick allele (ALT_CRICK), sample names.
- The original scripts used usearch for clustering the assembled or joined reads and creating a *de novo* reference. This was changed to vsearch^27^ (version 2.5.0) in epiGBS2.

## Access of the pipeline and lab protocol

The epiGBS2 lab protocol and pipeline documentation can be found in the supplemental material. The bioinformatics pipeline can be accessed on github (https://github.com/nioo-knaw/epiGBS2) and was deposited on zenodo (https://doi.org/10.5281/zenodo.3819996).

## Example Workflow and Output

The epiGBS2 lab procedure starts with extracted DNA, which is free of ethanol and secondary metabolites. For more details, please refer to the laboratory protocol in the supplemental materials. After executing a paired-end next generation sequencing run, the sequencing reads should be 5’-adapter trimmed but custom parts (UMI/ “Wobble”, barcode, control nucleotide and restriction site overhang) should remain. The reads of individual samples are still multiplexed, so you will receive two input files for the bioinformatics workflow in fastq format: Read 1 (forward reads, usually indicated by “R1” in the file name) and Read 2 (reverse reads, usually indicated by “R2” in the file name).

The following steps have to be taken to successfully run the workflow. For a more detailed description, please read the workflow documentation in the supplemental material.

1. Make sure that technical requirements are matched.
2. Copy the epiGBS2 pipeline from github (https://github.com/nioo-knaw/epiGBS2) or zenodo (https://doi.org/10.5281/zenodo.3819996).
3. Fill in the config file.
4. Prepare a barcode file.
5. Start the pipeline (make sure to use e.g. tmux if not working on a cluster, so results will not be lost, if you accidentally close the terminal).
6. Check the status of the pipeline regularly for errors.
7. After everything is finished, inspect the report.html and multiQC report.
8. Check output files as described in the documentation.

The SNP calls are summarized for each sample in a vcf format, from which you can e.g. plot SNP depth as shown in Figure 3A and perform downstream genetic analysis, such as genetic map construction, population genomics or phylogenetics. All predicted methylation sites are reported in the methylation.bed file and their genomic context (CHH, CG or CHG) is returned (Fig. 3B). Counts of total reads and methylated reads are recorded for each cytosine in each sample. These counts can be used to determine the number of samples covered at each position (Fig. 3C) and in the further downstream analysis to set a threshold to only keep positions with a minimum of covered samples. Total and methylated read counts are also needed to calculate methylation ratios (Fig 3D), on which differential methylation analysis can be based.

**Figure 3:**
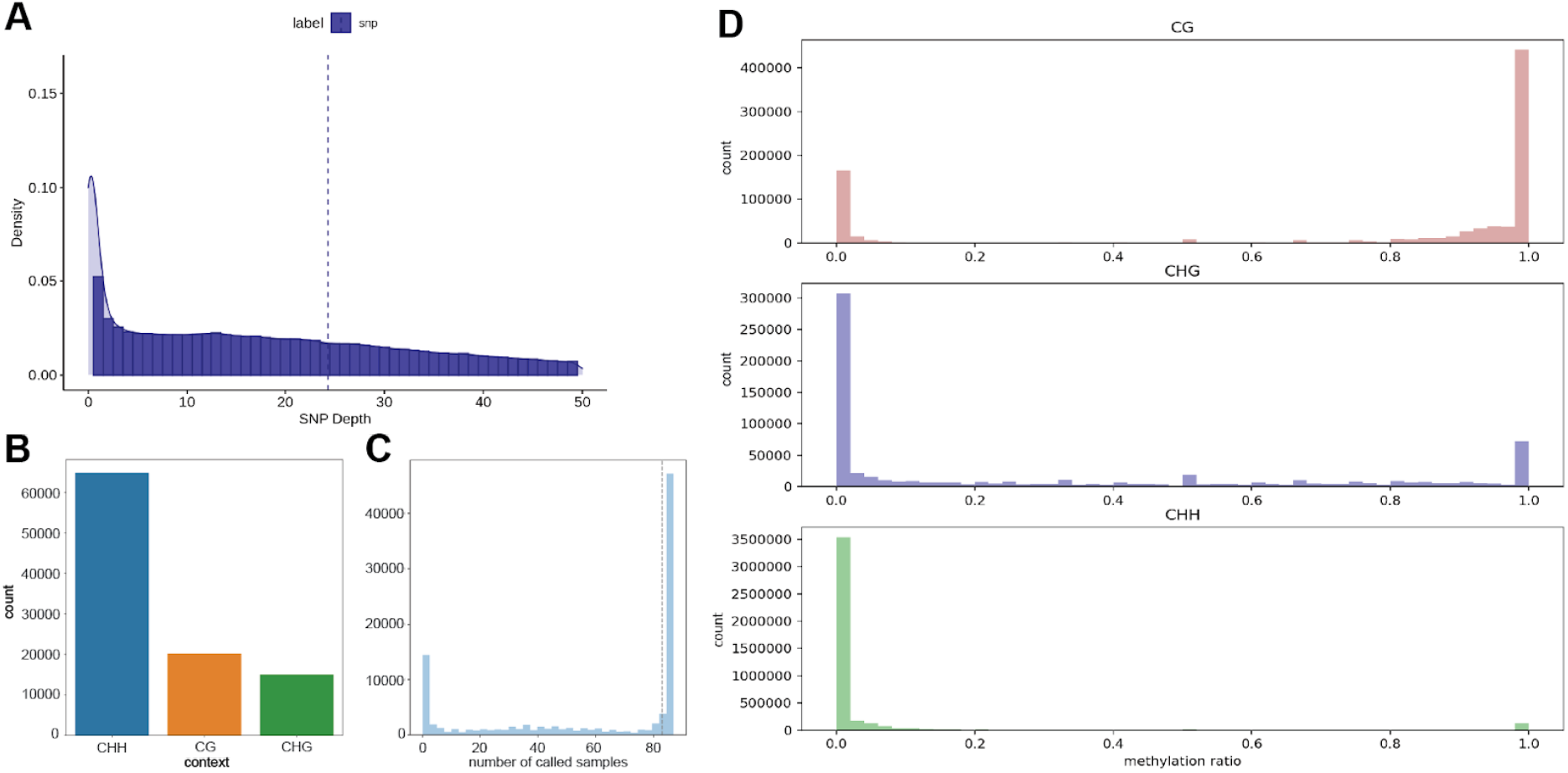
Example Output of epiGBS2. A) A file with SNPs is generated by the pipeline in.vcf format and for quality control SNP depth distribution can be plotted. Methylation information is stored in a.bed formatted file, from which main downstream analyses can be performed. B) The genomic sequence of the cytosine positions (CHH, CG or CHG) can be extracted and plotted. In plants, typically, the CHH context is most abundant. C) Per position the number of absent/present samples can be determined, plotted and used as a filter criterium to remove the least represented positions. D) Total and methylated read counts are used to calculate the methylation ratio, which can be used for downstream differential methylation analysis. Here, the methylation ratio is plotted in each genomic context. Our example is representative of the variation in methylation ratio that we usually observe in plants.

## Conclusion and Future Research

We presented here epiGBS2, consisting of a detailed description of our current laboratory protocol and a bioinformatics workflow that we use for epiGBS analysis. EpiGBS2 includes several major updates compared to the method as first published by van Gurp *et al.* (2016) and aims at the researcher with some basic experience in bioinformatics, e.g. working in a Linux environment. The updates were made in part to improve the method but also to make the method available in a more user-friendly and reproducible way to users working on a wide range of organisms. Where up to date, published papers using epiGBS have been mainly limited to plant species, the restriction enzyme flexibility of epiGBS2 does allow for its use in organisms, where RRBS used to be the preferred method.

Making epiGBS2 available allows others to use the updated methodology. However, we note that the development of epiGBS is work in progress and not all aspects of the methods are exhaustively benchmarked yet. Therefore, although the epiGBS2 pipeline is straightforward to execute and comes with some quality assessments of the analysis, the user should thoroughly check any obtained results. Some known but minor shortcomings of the bioinformatics procedures are the following: 1) During demultiplexing the presence of the expected RE overhangs are validated. This validation accepts one nucleotide mismatch to allow recognition of C-to-T converted RE overhang sequences after bisulfite treatment. If a mismatching nucleotide is identified (e.g. a T instead of a C), it is replaced with a C by the stacks code; this can effectively result in an unmethylated cytosine becoming labeled as methylated. One possible solution could be for a script to approve the remaining RE overhang and allow C/T conversions without replacing them. 2) The reference branch was built from scratch and currently exists for experimental purposes to facilitate epiGBS analysis based on an existing reference genome. When testing this reference branch we typically observe lower mapping percentages than for the *de novo* branch. This indicates that further benchmarking and parameter optimization for this reference branch are priorities for future work.

To further improve reliability of the bioinformatics workflow, future work should also include benchmarking of the mapping, SNP calling and methylation calling, beyond what has been done and presented in the 2016 paper. All these processes are largely based on custom scripts that have not yet been widely tested by the scientific community. Additionally, comparisons with existing similar pipelines, such as BsRADSeq^5^ for the *de novo* approach or Bismark^8^ for a reference-based analysis, can be performed to evaluate the relative performance of the presented workflow. Alternatively, if preferred, epiGBS2-generated data can also be aligned and analyzed using other existing methylation pipelines.

Regarding the laboratory protocol, future changes are expected in relation to the movement in sequencing platforms. The current protocols have been optimized for Illumina HiSeq systems, but sequencing agencies are currently changing to the NovaSeq systems. To allow pooling with libraries from other users on those high-throughput machines, adapters will have to be adjusted and a common Illumina barcode will have to be included. Another improvement could be made in achieving uniformity in sequencing output per sample. The epiGBS protocol prioritizes efficiency and cost-effectiveness in library preparation; however, in our hands, we observe considerable between-sample variability in read counts after sequencing. This could be improved by quantifying DNA amounts per individual sample with a qPCR step after adapter ligation. This might make the library preparation process more elaborate, but will provide more control over individual sample output. Alternatively, the sequencing performance of individual barcodes and barcode combinations could be benchmarked in more detail.

## Supporting information

pipeline documentation

Laboratory protocol

report with example data

## Acknowledgments

Bernice Sepers, Teresa Boquete, Nelia Luviano, Verónica Noé Ibañez, Cristian Peña, Veronika Laine, Felix Bartusch, Anupoma Niloya Troyee and Adam Nunn are acknowledged for critical questions, discussions and testing the pipeline. The contributions of MVA and SF were enabled by the European Training Network “EpiDiverse”, which received funding from the EU Horizon 2020 program under Marie Skłodowska-Curie grant agreement No 764965. MP and PV were funded by an Aspasia grant PV.

## Supplemental Material

### Protocols

- Laboratory protocol for library preparation
- Manual and documentation of the computational workflow in.html

### Tabels

- meta-analysis on the term BSseq on google scholar and in SRA

### Others

- example report file in.html

## References

1. Reyna-López, G. E., Simpson, J. & Ruiz-Herrera, J. Differences in DNA methylation patterns are detectable during the dimorphic transition of fungi by amplification of restriction polymorphisms. Mol. Gen. Genet. MGG 253, 703–710 (1997).

2. Suzuki, M. et al. Whole-genome bisulfite sequencing with improved accuracy and cost. Genome Res. 28, 1364–1371 (2018).

3. Meissner, A. et al. Reduced representation bisulfite sequencing for comparative high-resolution DNA methylation analysis. Nucleic Acids Res. 33, 5868–5877 (2005).

4. Gurp, T. P. van et al. epiGBS: reference-free reduced representation bisulfite sequencing. Nat. Methods 13, 322–324 (2016).

5. Trucchi, E. et al. BsRADseq: screening DNA methylation in natural populations of non-model species. Mol. Ecol. 25, 1697–1713 (2016).

6. Schield, D. R. et al. EpiRADseq: scalable analysis of genomewide patterns of methylation using next-generation sequencing. Methods Ecol. Evol. 7, 60–69 (2016).

7. Guo, W. et al. BS-Seeker2: a versatile aligning pipeline for bisulfite sequencing data. BMC Genomics 14, 774 (2013).

8. Bismark: a flexible aligner and methylation caller for Bisulfite-Seq applications | Bioinformatics | Oxford Academic. https://academic.oup.com/bioinformatics/article/27/11/1571/216956.

9. Kretzmer, H., Otto, C. & Hoffmann, S. BAT: Bisulfite Analysis Toolkit. F1000Research 6, 1490 (2017).

10. Richards, C. L. et al. Ecological plant epigenetics: Evidence from model and non-model species, and the way forward. bioRxiv 130708 (2017) doi:10.1101/130708.

11. Sepers, B. et al. Avian ecological epigenetics: pitfalls and promises. J. Ornithol. 160, 1183–1203 (2019).

12. Boquete, M. T., Wagemaker, N. C. A. M., Vergeer, P., Mounger, J. & Richards, C. L. Epigenetic Approaches in Non-Model Plants. in Plant Epigenetics and Epigenomics: Methods and Protocols (eds. Spillane, C. & McKeown, P.) 203–215 (Springer US, 2020). doi:10.1007/978-1-0716-0179-2_14.

13. Moorsel, S. J. van et al. Evidence for rapid evolution in a grassland biodiversity experiment. Mol. Ecol. 28, 4097–4117 (2019).

14. Di Tommaso, P. et al. Nextflow enables reproducible computational workflows. Nat. Biotechnol. 35, 316–319 (2017).

15. Köster, J. & Rahmann, S. Snakemake—a scalable bioinformatics workflow engine. Bioinformatics 28, 2520–2522 (2012).

16. Perkel, J. M. Workflow systems turn raw data into scientific knowledge. Nature http://www.nature.com/articles/d41586-019-02619-z (2019) doi:10.1038/d41586-019-02619-z.

17. Grüning, B. et al. Bioconda: sustainable and comprehensive software distribution for the life sciences. Nat. Methods 15, 475–476 (2018).

18. Stacks 2: Analytical Methods for Paired-end Sequencing Improve RADseq-based Population Genomics | bioRxiv. https://www.biorxiv.org/content/10.1101/615385v1.

19. Catchen, J. M., Amores, A., Hohenlohe, P., Cresko, W. & Postlethwait, J. H. Stacks: Building and Genotyping Loci De Novo From Short-Read Sequences. G3 Genes Genomes Genet. 1, 171–182 (2011).

20. Catchen, J., Hohenlohe, P. A., Bassham, S., Amores, A. & Cresko, W. A. Stacks: an analysis tool set for population genomics. Mol. Ecol. 22, 3124–3140 (2013).

21. Pedersen, B. S., Eyring, K., De, S., Yang, I. V. & Schwartz, D. A. Fast and accurate alignment of long bisulfite-seq reads. ArXiv14011129 Q-Bio (2014).

22. Dobin, A. et al. STAR: ultrafast universal RNA-seq aligner. Bioinformatics 29, 15–21 (2013).

23. Garrison, E. & Marth, G. Haplotype-based variant detection from short-read sequencing. ArXiv12073907 Q-Bio (2012).

24. Li, H. et al. The Sequence Alignment/Map format and SAMtools. Bioinformatics 25, 2078–2079 (2009).

25. Ewels, P., Magnusson, M., Lundin, S. & Käller, M. MultiQC: summarize analysis results for multiple tools and samples in a single report. Bioinformatics 32, 3047–3048 (2016).

26. Martin, M. Cutadapt removes adapter sequences from high-throughput sequencing reads. EMBnet.journal 17, 10–12 (2011).

27. Rognes, T., Flouri, T., Nichols, B., Quince, C. & Mahé, F. VSEARCH: a versatile open source tool for metagenomics. PeerJ 4, (2016).

28. Andrews, Simon. FastQC: a quality control tool for high throughput sequence data. Available online at: http://www.bioinformatics.babraham.ac.uk/projects/fastqc. (2010).

29. Zhang, J., Kobert, K., Flouri, T. & Stamatakis, A. PEAR: a fast and accurate Illumina Paired-End reAd mergeR. Bioinformatics 30, 614–620 (2014).

