## Supplementary material for "epiGBS2: an improved protocol and automated snakemake workflow for highly multiplexed reduced representation bisulfite sequencing": pipeline documentation: epiGBS-documentation.html

README.utf8.md


### Manual epiGBS2

- Prerequisites for running the pipeline
- Preparation to run the pipeline
- Start the pipeline
- Explanation of files in the output directory
- When not to run the pipeline?
- Quality control or “How to discover errors?”
- Example Config Files
- List of used software and references

#### Prerequisites of bioinformatics skills and infrastructure and wetlab preparations for running the pipeline

- A basic knowledge of Linux:
  - Knowledge, about how to work with files and directories (cd, ls, nano)
  - Being able to execute commands (git, conda, snakemake)
- Knowledge about the statistical analysis of e.g. differential methylation. This is not included in the pipeline.
- Linux server:
  - e.g. Ubuntu 16.04
  - Sudo rights not necessary
  - Miniconda installed or
    - Download with `wget` https://docs.conda.io/en/latest/miniconda.html
    - Installing like described here, can be installed in user’s home without sudo
      - `bash Miniconda3-latest-Linux-x86_64.sh`
- Adapters include:
  - Control nucleotide
  - Wobble (running without is also possible)
- Sequencing:
  - paired-end Illumina-sequencing
  - adjust output (number of reads) according to your genome size and expected number of fragments (e.g. based on an *in silico* digest)
- Readfiles:
  - un-demultiplexed but standard Illumina adapter trimmed (usually already done by sequencing agency)
- No reference genome required

#### Preparation to run the pipeline

- Make a conda environment for snakemake if snakemake is not installed globally on the server. You do not need administrator rights to do this but conda has to be installed (see Prerequisites for running the pipeline).
  - `conda create -n snake snakemake=5.4.5`
  - `conda activate snake`
- Make a copy of the pipeline
  - `git clone https://github.com/nioo-knaw/epiGBS2.git`
- Enter the created directory:
  - `cd epiGBS2`
- Open and adjust the config file: **All paths are full paths, no relative paths allowed.** For examples, please see Example Config Files
  - `nano config.yaml`
  - output\_dir: Path of directory to store all output files and directory. Path will be created by the pipeline and should not pre-exist. E.g. if the path of the cloned directory is `/fleurg/projects/epiGBS2`, then use `/fleurg/projects/epiGBS2/output`.
  - input\_dir: Path of directory containing raw data (e.g. fastq.gz, or fq.gz) and barcode file
  - read1: Name of the read file containing forward (R1) reads
  - read2: Name of the read file containing forward (R2) reads
  - cycles: read length
  - barcodes: Filename of the barcode file (do not include the path!)
  - tmpdir: Path to the directory where temporary files will be stored. For most systems this will be /tmp
  - threads: number of available computing threads on your system
  - mode: choice between “denovo” or “reference”
  - ref\_dir: Only needed for reference-based analysis. Path to directory containing a reference genome
  - Genome: Only needed for reference-based analysis. Name of the genome file (prefix .fa)
  - Param\_denovo: Optional and only relevant for analysis in denovo mode. To run on default choose "" or “default”, otherwise enter a number
    - Identity: percentage of sequence identity in the last clustering step, in decimal number e.g. for 90% identity write 0.90, default: “0.95”
    - Min-depth: minimal cluster depth in the first clustering step to include a cluster, default 0
    - Max-depth: maximal cluster depth in the first clustering step to include a cluster, default 0
  - param\_SNPcalling:
    - max-depth: At a position, read maximally INT reads per input file, default 10000000, check if mpileup automatically reduced the maximal depth even more. You might have to re-run your analysis with a lower max-depth value to avoid this.
    - min-MQ: Minimum mapping quality for an alignment to be used, default 0
    - min-BQ: Minimum base quality for a base to be considered, default 15
- Make a barcode file: The barcode file is tab-delimited and contains at least the following columns: Flowcell, Lane, Barcode\_R1, Barcode\_R2, Sample, ENZ\_R1, ENZ\_R2, Wobble\_R1, Wobble\_R2. All other fields are optional. Make sure that the sample name does not only contain numbers but also letters. If you prepare the barcode file in Excel, make sure that no `^M` are present after uploading the file to the Linux server. You can check this by opening the barcode file with `cat -A barcode.tsv` on a Linux server. You can remove the `^M` with `sed -e "s/^M//" filename > newfilename`. To enter ^M, type CTRL-V, then CTRL-M. That is, hold down the CTRL key then press V and M in succession.

The Flowcellname can be found in the fastq headers of the read file, e.g. `@ST-E00317:403:H53KHCCXY:5:1101:5660:1309 1:N:0:NCAATCAC` translates to `@ST-E00317:403:FLOWCELL:LANE-NUMBER:1101:5660:1309 1:N:0:NCAATCAC`. ENZ\_R1/2 expects the names of the restriction enzymes and Wobble\_R1/2 is the length of the unique molecular identifier (“Wobble”) sequence (usually 3).

```
# barcodes.tsv
Flowcell        Lane    Barcode_R1      Barcode_R2      Sample  history Country PlateName       Row     Column  ENZ_R1  ENZ_R2  Wobble_R1       Wobble_R2       Species
H53KHCCXY       5       AACT    CCAG    BUXTON_178      C       BUXTON  BUXTON_WUR_AseI_NsiI_final_run1 1       2       AseI    NsiI    3       3       Scabiosa columbaria
H53KHCCXY       5       CCTA    CCAG    WUR_178 C       WUR     BUXTON_WUR_AseI_NsiI_final_run1 2       2       AseI    NsiI    3       3       Scabiosa columbaria
H53KHCCXY       5       TTAC    CCAG    BUXTON_169      C       BUXTON  BUXTON_WUR_AseI_NsiI_final_run1 3       2       AseI    NsiI    3       3       Scabiosa columbaria
H53KHCCXY       5       AGGC    CCAG    WUR_169 C       WUR     BUXTON_WUR_AseI_NsiI_final_run1 4       2       AseI    NsiI    3       3       Scabiosa columbaria
H53KHCCXY       5       GAAGA   CCAG    BUXTON_175      SD      BUXTON  BUXTON_WUR_AseI_NsiI_final_run1 5       2       AseI    NsiI    3       3       Scabiosa columbaria
H53KHCCXY       5       CCTTC   CCAG    WUR_175 SD      WUR     BUXTON_WUR_AseI_NsiI_final_run1 6       2       AseI    NsiI    3       3       Scabiosa columbaria
```

#### Start the pipeline

**Running the pipeline with `--use-conda` is important!**

- Dry-run:
  - `snakemake -n --use-conda`
- everythin green? Then…
- run the pipeline: `snakemake -j <threads> -p --use-conda` Replace  by number of CPU’s to use on your server, e.g. `snakemake -j 12 -p --use-conda`

#### Explanation of files in the output directory

It follows a description of all output files. Files that are important for downstream analysis are highlighted in bold. Files or Directories in italics are specific for the de-novo and reference branch respectively.

- **report.html**: A report summarizing all stats from the epiGBS analysis. Absolutely crucial to determine, whether your analysis ran successfully or not. However, this file gives only a first impression and further analysis is necessary to confirm the quality of your analysis.
- **multiQC\_report.html**: A report summarizing QC stats for the input data. Can be opened in any webbrowser. This file can be very large. Hide parts of the files to make loading easier, e.g. filter out all files containing “rem”
- output\_demultiplex:
  - barcode\_stacks.tsv: barcode file converted to the required stacks format
  - clone: Directory containing read files from which PCR duplicates were removed
  - clone-stacks: Directory containing demultiplexed and Watson-Crick separated read files. File names start with sample names. Files with *rem* in the name contain reads that failed the RAD-tag check.
    - process\_radtags.clone.log: Log file containing stats from demultiplexing, also contains information about eventually discovered *de novo* barcodes
  - crick\_R1.fq.gz: All filtered and demultiplexed crick forward reads, input for following steps
  - crick\_R2.fq.gz: All filtered and demultiplexed crick reverse reads, input for the following steps
  - demultiplex.log: Log file containing stats from clone-removal
  - Watson\_R1.fq.gz: All filtered and demultiplexed watson forward reads, input for the following steps
  - Watson\_R2.fq.gz: All filtered and demultiplexed watson reverse reads, input for the following steps
- *output\_denovo*: Containing all files created during de-novo reference creation. Skipped, if running in reference-mode.
  - Assembled.fq.gz: All read pairs (forward/R1 and reverse/R2), that were assembled/merged using Pear
  - Unassembled.R1.fq.gz: All forward/R1 reads that could not be assembled/merged
  - Unassembled.R2.fq.gz: All reverse/R2 reads that could not be assembled/merged
  - consensus.fa: sequence file. outputfile from second clustering step, in which binary watson and crick reads are matched, and after reference reconstruction
  - consensus\_cluster.fa: De novo reference sequence file. Outputfile from third clustering step, in which sequences from previous steps were clustered based on identity
  - **consensus\_cluster.renamed.fa**: sequences are identical to consensus\_cluster.fa but fasta names were renamed. Input for mapping.
  - consensus\_cluster.renamed.fa.fai: Indexed *de novo* reference sequences.
  - make\_reference.log: Log file containing some stats about the clustering.
- mapping:
  - STAR\_{joined,merged}\_{crick,watson}: Directory, which contains the (indexed) Crick/Watson-reference converted to a three-letter alphabet. If working with a reference genome, differentiation between joined/merged absent.
  - {crick, watson}.bam: Alignment file of crick or watson reads against the reference, created by STAR
  - {crick,watson}.bam.bai: indexed alignment files, used as input for samtools
  - {crick,watson}.vcf.gz: created by samtools, contains all variant positions in {crick,watson} reads against the reference for each sample
  - merged.tsv.gz: created by a custom script that merges crick.vcf.gz and watson.vcf.gz
  - **snp.vcf.gz**: created by a custom script that extracts all SNPs from merged.tsv.gz. You can read this file using zcat or after unzipping it using gunzip
  - header.sam: File containing the sam header
  - mapping\_variantcalling.log: Contains the mapping statistics
  - **methylation.bed**: created by a custom script that extracts all methylated positions for each sample
  - heatmap.igv: Input for IGV (https://software.broadinstitute.org/software/igv/) to visualize methylated positions in the genome
  - {Crick,Watson}\_{joined,merged}Log.final.out:
- *trimmed*: Directory containing adapter trimmed reads and their fastqc output files. For running in reference-mode only.
- multiQC\_report\_data: Directory containing all log and intermediary files that are created by MultiQC
- fastqc: Directory containing individual FastQC reports
- log: Directory containing log-files

#### When not to run the pipeline?

- If you want to determine methylation in restriction enzyme overhang. The original sequence will be replaced by the expected overhang during demultiplexing if a mismatch between expected and actual overhang sequence occurs. Hence, C-T conversions are replaced by a C and methylation would be artifically set to 100%.
- The reference branch is in an experimental stage. One observed drawback is a low mapping percentage (20-30 %) but it might depend from the reference genome and organism.
- SNP and methylation calling are not benchmarked.

#### Quality control or “How to discover errors?”

Recommendation: Run fastq-screen in bisulphite mode on raw data to determine sources of contamination (e.g. by sharing a lane with other customers, human DNA, phiX, vectors and adapters). https://www.bioinformatics.babraham.ac.uk/projects/fastq\_screen/\_build/html/index.html and https://www.bioinformatics.babraham.ac.uk/projects/fastq\_screen/fastq\_screen\_documentation.html

##### MultiQC report:

- MultiQC wraps the FastQC, mapping and trimming (for reference-branch only) report of all samples and gives you the possibility to compare the quality plots of all or specific samples directly with each other. All functions of MultiQC are explained in a tutorial video that is provided as a link in the report.
- you can consider the FastQC documentation to understand most of the FastQC plots: https://www.bioinformatics.babraham.ac.uk/projects/fastqc/Help/3%20Analysis%20Modules/. However, epiGBS libraries have the following specific characteristics that will be reflected in the quality reports. You can also compare this with the RRBS example from the FastQC website: https://www.bioinformatics.babraham.ac.uk/projects/fastqc/RRBS\_fastqc.html
  - The sequence duplication levels will be higher than average because restriction enzymes were used to create the libraries versus random shearing for e.g. whole genome sequencing.
  - The per base sequence content of the first nucleotides will re-construct the overhang of the used restriction enzyme after demultiplexing the reads. The “C” content is usually low and “T” is high.
  - The 3’-end adapter content is usually high

##### EpiGBS-report:

- Number of clones: Check the number and distributions of clones. The dominant peak should be at a number of clones = 1.
- Demultiplexing: The number of reads per sample should be comparable and agree with the expected numbers. The amount of Watson and Crick reads for a specific sample should be similar, too. The optimal number of reads depends on the expected number of fragments (> 10 reads per fragment) and required coverage.
- *De novo* reference:
  - Check the number of assembled reads. Usually the percentage of assembled reads should be higher than unassembled reads. This depends from the size of the fragments that you expect. If the fragment size is greater than twice the chosen read site, the amount of assembled reads will be small.
  - The number of consensus clusters (third clustering step) should be comparable with the number of fragments that you expected, e.g. by performing an in silico digest using R packages like SimRAD.
- Mapping:
  - The mapping percentage for all reads should be higher than 40%. In general, we observe a lower mapping percentage for assembled than for joined reads.
- SNP calling:
  - The amount of detected SNPs will depend from the used species and the diversity of samples. The SNP depth should be greater than 10 for reliable calling.
- Methylation calling:
  - The amount of methylated cytosines and their context will depend from the used species and used restriction enzymes. In general, the depth should be greater than 10 for reliable calling.

#### Fix errors

##### Clone percentage

###### Problem:

I have a very high percentage of clone reads

###### Fix:

- Check the length of the Wobble in your adapter sequence and the number in the barcode file.
- Check the quality and quantity of input DNA during the wetlab procedure.

##### Demultiplexing

###### Problem:

One or more samples have small amounts of recovered reads or read numbers differ a lot between different samples.

###### Fix:

- Check the labwork (e.g. the used barcodes and your pipetting scheme)
- Check the barcode file based on the wetlab-scheme
- The Barcode log file (output.dir/output-demultiplex/clone-stacks/process\_radtags.clone.log) contains denovo discovered barcodes. Do you find a complete set of Watson and Crick, reverse and forward? Then check your barcode file and experimental design file again. Do you find a lot of barcode-sets, where both R1 and R2 barcode end on “C”? Then bisulfite conversion might be low in your experiment.
- Check DNA quantification and quality of the missing samples
- RAD-tag: Pipeline checks for the presence of the restriction enzyme overhang (RAD-tag). If this sequence contains unmethylated “Cs”, they will be converted to “Ts”. The RAD-tag check allows one nucleotide mismatch. Depending from the chosen enzyme combination, two or more mismatches should be allowed. Possible fix is to switch off RAD-tag check by opening demultiplex/barcode\_stacks.py and change line XX from cmd += “-r -D –inline\_inline –barcode\_dist\_2 0” to cmd += “-r -D –inline\_inline –barcode\_dist\_2 0 –disable\_rad\_check”
- Check the .rem. files in output.dir/output-demultiplex/clone-stacks/ in the QC. Do they show a common sequence in the beginning of the read different from your expected RAD-tag?
- check for barcode bias. From GBS experiment it is known that some barcodes perform better than others

###### Problem:

The coverage is too low in the methylation bed file and after filtering on coverage (>10) only few positions remain.

###### Fix:

- The required amount of reads will depend from the expected number of DNA fragments. You can calculate this for your (or a related) species by using R packages like SimRAD (https://cran.r-project.org/package=SimRAD)
- sequence more
- reduce the genome representation by using a low cutting restriction enzyme

#### Example Config Files

##### De novo

```
# path to output directory
output_dir: "/fleurg/projects/epiGBS2/output"

# input directory where raw reads are
input_dir       : "/fleurg/projects/epiGBS2/data"

# name of sequence read files
Read1 : "epiGBS_1.fq.gz"
Read2 : "epiGBS_2.fq.gz"

# number of sequencing cycles (the same as read length in Illumina sequencing)
cycles        : 150

# barcode file(barcode file should be kept inside input directory) and enzymes will be included in barcode file
barcodes: "barcodes.tsv"

# the pipeline produces some temporary files. Please indicate the tmp location on your server (in most cases /tmp)
tmpdir        : "/tmp"

# some of the steps can be run in parallel. Please set the number of available computing threads on your system
threads: "12"

# mode of running pipeline (set denovo or reference)
mode: "denovo"

# genome directory (leaave it blank in denovo mode)
ref_dir: ""

# genome name (leaave it blank in denovo mode)
genome: ""

# advanced users have the possibility to change different parameter, leave them blank or write "default" to run them in default mode

# parameters in the denovo reference creation:
# identity: percentage of sequence identity in the last clustering step, in decimal number e.g. for 90% identity write 0.90, default 0.95
# min-depth: minimal cluster depth in the first clustering step to include a cluster, default 0
# max-depth: maximal cluster depth in the first clustering step to include a cluster, default 0
param_denovo:
  identity: ""
  min-depth: ""
  max-depth: ""

# parameters in the mpileup step (variant callin)
# max-depth: At a position, read maximally INT reads per input file, default 10000000, check if mpileup automatically reduced the maximal depth even more. You might have to re-run your analysis with a lower max-depth value to avoid this.
# min-MQ: Minimum mapping quality for an alignment to be used, default 0
# min-BQ: Minimum base quality for a base to be considered, default 15

param_SNPcalling:
  max-depth: ""
  min-MQ: ""
  min-BQ: ""
```

##### Reference

```
# path to output directory
output_dir: "/fleurg/projects/epiGBS2-ref/output"

# input directory where raw reads are
input_dir       : "/fleurg/projects/epiGBS2-ref/data"

# name of sequence read files
Read1 : "epiGBS_1.fq.gz"
Read2 : "epiGBS_2.fq.gz"

# number of sequencing cycles (the same as read length in Illumina sequencing)
cycles        : 150

# barcode file(barcode file should be kept inside input directory) and enzymes will be included in barcode file
barcodes: "barcodes.tsv"

# the pipeline produces some temporary files. Please indicate the tmp location on your server (in most cases /tmp)
tmpdir        : "/tmp"

# some of the steps can be run in parallel. Please set the number of available computing threads on your system
threads: "12"

# mode of running pipeline (set denovo or reference)
mode: "reference"

# genome directory (leaave it blank in denovo mode)
ref_dir: "/fleurg/projects/epiGBS2-ref/data/ref"

# genome name (leave it blank in denovo mode)
genome: "reference.fa"

# advanced users have the possibility to change different parameter, leave them blank or write "default" to run them in default mode

# parameters in the denovo reference creation:
# identity: percentage of sequence identity in the last clustering step, in decimal number e.g. for 90% identity write 0.90, default 0.95
# min-depth: minimal cluster depth in the first clustering step to include a cluster, default 0
# max-depth: maximal cluster depth in the first clustering step to include a cluster, default 0
param_denovo:
  identity: ""
  min-depth: ""
  max-depth: ""

# parameters in the mpileup step (variant callin)
# max-depth: At a position, read maximally INT reads per input file, default 10000000, check if mpileup automatically reduced the maximal depth even more. You might have to re-run your analysis with a lower max-depth value to avoid this.
# min-MQ: Minimum mapping quality for an alignment to be used, default 0
# min-BQ: Minimum base quality for a base to be considered, default 15

param_SNPcalling:
  max-depth: ""
  min-MQ: ""
  min-BQ: ""
```

#### List of used software and references

##### Software

- epiGBS2
- Snakemake 5.4.5
- Conda
- Stacks
- Python 3.7
- R + R package to render Rmd
- FastQC
- Trim-galore
- Pear
- Seqtk
- STAR
- vsearch
- Samtools

##### References

1. Manuscript on bioRxiv
2. Köster, J. & Rahmann, S. Snakemake-a scalable bioinformatics workflow engine. Bioinformatics 28, 2520-2522 (2012).
3. Grüning, B. et al. Bioconda: sustainable and comprehensive software distribution for the life sciences. Nat. Methods 15, 475-476 (2018).
4. Catchen, J., Hohenlohe, P. A., Bassham, S., Amores, A. & Cresko, W. A. Stacks: an analysis tool set for population genomics. Mol. Ecol. 22, 3124-3140 (2013).
5. Catchen, J. M., Amores, A., Hohenlohe, P., Cresko, W. & Postlethwait, J. H. Stacks: Building and Genotyping Loci De Novo From Short-Read Sequences. G3 Genes Genomes Genet. 1, 171-182 (2011).
6. Stacks 2: Analytical Methods for Paired-end Sequencing Improve RADseq-based Population Genomics | bioRxiv. Available at: https://www.biorxiv.org/content/10.1101/615385v1. (Accessed: 27th August 2019)
7. Andrews, Simon. FastQC: a quality control tool for high throughput sequence data. Available online at: http://www.bioinformatics.babraham.ac.uk/projects/fastqc. (2010).
8. Zhang, J., Kobert, K., Flouri, T. & Stamatakis, A. PEAR: a fast and accurate Illumina Paired-End reAd mergeR. Bioinformatics 30, 614-620 (2014).
9. Dobin, A. et al. STAR: ultrafast universal RNA-seq aligner. Bioinformatics 29, 15-21 (2013).
10. Rognes, T., Flouri, T., Nichols, B., Quince, C. & Mahé, F. VSEARCH: a versatile open source tool for metagenomics. PeerJ 4, (2016).
11. Lepais, O. & Weir, J. T. SimRAD: an R package for simulation-based prediction of the number of loci expected in RADseq and similar genotyping by sequencing approaches. Mol. Ecol. Resour. 14, 1314-1321 (2014).
