## Supplementary material for "epiGBS2: an improved protocol and automated snakemake workflow for highly multiplexed reduced representation bisulfite sequencing": Laboratory protocol

### EpiGBS2 Laboratory User Manual

Table of Contents

**1. Reagents, Consumables and Equipment** **2**
1.1 Reagents 2
1.2 Consumables 3
1.3 Equipment 3
1.4 Oligo-nucleotide sequences 4
**2. Protocol description** **4**
2.1 Introduction 4
2.2 Starting Biological Material 5
2.2.1. DNA isolation from plant tissue 5
2.2.2 DNA isolation from bird 5
2.3 Adapter Design 6
**3. epiGBS Protocol** **7**
3.1 DNA digestion 7
3.2 Annealing Adapters 7
3.3 Adapter Ligation 8
3.4 Multiplexing 9
3.5 Concentrating and Cleaning 9
3.6 0.8X SPRI size-selection 10
3.7 Nick Repair 11
3.8 Test PCR (Optional) 12
3.9 Bisulfite Conversion (BS) 13
3.10 Library amplification PCR 13
3.11 Library Cleaning 15
3.12 0.8X SPRI size-selection 15
3.13 Library Quality Check 17
3.14 Sequencing 18

### 1. Reagents, Consumables and Equipment

#### 1.1 Reagents

| **Reagent** | **Brand** | **Supplier** | **Catalogue Number** | **Amount** |
|---|---|---|---|---|
| **Restriction Enzymes **** | | | | |
| Fast NsiI | NEB | Bioke | R0127L | 5000 units |
| AseI | NEB | Bioke | R0526L | 10.000 units |
| MspI | NEB | Bioke | R0106S | 5000 units |
| Adapter Ligation | | | | |
| T4 DNA ligase (2.000.000 units/ml) | NEB | Bioke | M0202M | 100.000 units |
| **Concentrating and Cleaning** | | | | |
| NucleoSpin Gel and PCR Cleanup Kit | Marchery Nagel | Bioke | 740609.250 | 250 preps |
| NucleoSpin buffer BE (Tris-buffer) | Marchery Nagel | Bioke | 740306.100 | 125 ml |
| **0.8 x SPRI size selection** | | | | |
| Agencourt AMPure XP | Beckman Coulter | Beckman Coulter | A63881 | 60 ml |
| **Nick Repair** | | | | |
| 5-Methylcytosine dNTP Mix [10mM] | Zymo | BaseClear | D1030 | 2.5 µmol/250 µl |
| DNA Polymerase I | NEB | Bioke | M0209S | 500 units |
| NEB2 Buffer | NEB | Bioke | B7002S | 5 ml |
| **Bisulfite Conversion** | | | | |
| EZ DNA Methylation-Lightning Kit | Zymo | BaseClear | D5030 | 50 rxns |
| **Library Amplification** | | | | |
| KAPA HiFi Hot Start* Readymix | Roche diagnostics | Roche sequencing | KK2601 | 1.25 ml |

| | | | | |
|---|---|---|---|---|
| KAPA HiFI HS Uracil+ ReadyMix | Roche diagnostics | Roche sequencing | KK2801 | 1.25 ml |
| KAPA Library Quantification Kit | Roche diagnostics | Roche sequencing | KK4844 | 500 rxn |
| Agilent High Sensitivity DNA Kit | Agilent Technologies | Agilent genomics | DNF-474 | 500 samples |
| Qubit dsDNA HS Assay Kit | ThermoFisher Scientific | ThermoFisher Scientific | Q32854 | 500 assays |

** Enzymes choice can be modified depending on species of interest. It is recommended that you choose Restriction enzymes that are methylation insensitive and that are minimum 4 bp cutters.

#### 1.2 Consumables*

Eppendorf PCR tubes (0.5 ml)
Eppendorf DNA LoBind tubes (2.0 ml)
Eppendorf standard reaction tubes (1.5 ml)
Personal protection equipment (lab coat, gloves, goggles)

*Consumables which were used and tested when setting up the epiGBS2 laboratory protocol in the NIOO-KNAW molecular lab.

#### 1.3 Equipment **

Thermal heating-block
Incubated microplate shaker, VWR
Manual pipettes, Gilson
Eppendorf 5424R Centrifuge
Invitrogen Qubit 3.0 fluorometer
Biorad C1000 Touch Thermal cycler
Magnetic porter for 2 ml reaction tubes
Biorad CFX95 Touch Real time PCR detection system
Advanced Analytical Fragment Analyzer (Capillary electrophoresis)
Qiagen Tissuelizer II

** Equipment brands are those found in the NIOO-KNAW's molecular lab and have been tested to work with the epiGBS2 laboratory protocol.

#### 1.4 Oligo-nucleotide sequences

Standard Illumina paired end primers were used as shown in Table 1. Adapters were customized as described in 2.3 and consist of the standard illumina sequence, the unique molecular identifier or so called "Wobble", a barcode, the control nucleotide and the complement of the restriction enzyme overhang.

| Name | Sequence | comment | company |
|---|---|---|---|
| **Primer A** | 5'- AAT GAT ACG GCG ACC ACC GAG ATC T**AC ACT CTT TCC CTA CAC GAC GCT CTT CCG ATC T** - 3' | 100 nmol | IDT |
| **Primer B** | 5'- CAA GCA GAA GAC GGC ATA CGA GAT CGG T**CT CGG CAT TCC TGC TGA ACC GCT CTT CCG ATC T** - 3' | 100 nmol | IDT |
| **Adapter BA-I** | 5'- AXA XTX TTT XXX TAX AXG AXG XTX TTX XGA TXT **ZZZ AAXT C** -3' | X = 5mC, dephosph. | ALPHA DNA |
| **Adapter BA-II** | 5'- **TA G AGTT NNN** AGA TCG GAA GAG CGT CTG TAG GGA AAG AGT GT - 3' | dephosph. | ALPHA DNA |
| **Adapter CO-I** | 5'- XTX GGX ATT XXT GXT GAA XXG XTX TTX XGA TXT **ZZZ AAXT C TGXA** -3' | X = 5mC, dephosph. | ALPHA DNA |
| **Adapter CO-II** | 5'- **G AGTT NNN** AGA TCG GAA GAG CGG TTC AGC AGG AAT GCC GAG -3' | desphosph. | ALPHA DNA |

Tabel 1: oligonucleotide sequences used in this protocol. Bold: primer sequence that aligns to the adapter sequence, yellow: unique molecular identifier, "Wobble", orange: barcode, blue: control nucleotide, grey: complement to restriction enzyme sequence, desposph. : dephosphorylated

### 2. Protocol description

#### 2.1 Introduction

This epiGBS2 bench-ready laboratory protocol describes the library preparation for reduced representation bisulfite sequencing for various different species, including plants, vertebrates (*Parus major*) and invertebrates (*Biomphalaria glabrata*). Overall, this protocol follows Boquete *et al.* (2020) with the addition of describing how to design the customized adapters.

DNA is isolated individually from each sample and has to be ethanol-free and free of secondary metabolites (see point 3.1). DNA of individual samples is at first digested with a specific Restriction enzyme (RE) combination (see point 1.1 for commonly used REs and Figure 1 for a schematic overview of the protocol). After RE digestion, the obtained DNA fragments in each sample are ligated to customized adapters. These adapters include unique barcode combinations (see point 1.4 and 2.3) allowing for multiple samples to be multiplexed into a single sequencing library. Currently, 96 unique barcode combinations are available. However, the number of samples multiplexed together in the same library should be adapted depending on the targeted coverage, estimated read amount and genome size of the species.

Multiplexed samples are then concentrated and fragments sized 60 bp and smaller, which is the estimated size of adapter dimers, are removed. Next, 0.8x SPRI beads size selection is used to select for fragments of 150 bp and higher. Nick repair is then performed to 1) ligate the non-phosphorylated adapter ends and 2) replace non-methylated cytosines in the adapters by methylated cytosines. Subsequently, the selected ligated fragments are bisulfite treated, converting non-methylated Cytosines into Uraciles. Fragments are PCR amplified, cleaned and again 0.8X SPRI size selected to produce the final epiGBS library. The quantity and quality of the sequencing library is determined by qPCR quantification and on a fragment analyser.

#### 2.2 Starting Biological Material

Extracted DNA, free of ethanol and secondary metabolites, is needed to start the epiGBS2 laboratory protocol. Ideally, the DNA amount should range between 200 - 400 ng, but can be decreased to quantities as low as 30 ng. It is recommended to start the epiGBS2 protocol with 30 µl of DNA.

This laboratory protocol has been designed to work with a minimum of 12 and a maximum of 96 samples. It is recommended to work with a multiple of 12 samples (12, 24, 36, 48, 60, 72, 84 or 96 samples) to meet the requirements of our optimized setup.

##### 2.2.1. DNA isolation from plant tissue:

The Marchery Nagel NucleoSpin Plant II extraction kit was tested in several plant species and is compatible with the epiGBS2 laboratory protocol. DNA yield varied between 23 and 230 ng/µl depending on the species and overall quality of the DNA. In brief, the prefered homogenization method is through grinding, using VA steel beads and a tissue lyzer. The Marchery Nagel NucleoSpin Plant II extraction kit comes with two different lysis buffers. Both buffers have been tested and are compatible with the epiGBS2 laboratory protocol. However, if working with a new species, it is recommended that you test both lysis buffers before starting, as the efficiency of the lysis buffers can be species dependent.

##### 2.2.2 DNA isolation from bird:

As a starting material, 2 to 5 µl blood from *Parus major* (Great tit) was used and routinely DNA isolated with the FavorPrep™ 96-Well Genomic DNA Extraction Kit (Favorgen).

#### 2.3 Adapter Design

The epiGBS2 adapters consist of 4 customized, separately ordered oligonucleotides (see 1.4) that are annealed to two hemi-methylated adapters BA and CO (Fig. 1 and 3.2). All epiGBS2 adapters consist of a unique molecular identifier or so called “wobble sequence” followed by a barcode, the control nucleotide (a non-methylated cytosine) and the complement sequence of the restriction enzyme overhang sequence. Note that when designing new adapters, the barcode sequence and the complement of the restriction enzyme have to be adjusted. The choice of barcodes is crucial for the demultiplexing of individual samples computationally. It is recommended that the barcodes differ from one another by a minimum of three mutational base pairs (bp). R1/BA adapter barcodes should vary between 4 to 8 bp in length while the R2/CO barcode should be 4 bp long. In Table 2 multiple examples for BA and CO Barcodes are presented. Please note, if different RE combinations are used other than those suggested in this protocol, it is important to exchange the overhang sequence accordingly. Furthermore, it is important to ensure that the control nucleotide does not complement the restriction enzyme recognition site as this might lead to unintended digestion of ligated fragments.

| **BA Barcode** |
|---|
| AACT |
| CCTA |
| TTAC |
| AGGC |
| GAAGA |
| CCTTC |
| TTCAA |
| GCGGC |
| AGATGC |
| CATAGC |
| TTCGAC |
| ATGCGC |

| **CO Barcode** |
|---|
| AACT |
| CCAG |
| TTGA |
| GGTC |
| ACTA |
| CAGC |
| TGAT |
| GTCG |
| ATAC |
| CGCA |
| TATG |
| GCGT |
| GTAC |
| AGCT |

Table 2: List of possible barcode choices for both BA barcodes and CO barcodes. epiGBS was optimized to support 96 different barcode combinations in one sequencing library depending on which BA barcode is combined with which CO barcode.

### 3. epiGBS Protocol

#### 3.1 DNA digestion

1. In a reaction tube prepare a digestion master mix with the FastDigest buffer, Restriction enzymes (RE) and MiliQ water. Enzymes should be added last to the digestion mix.

   30 µl DNA
   -------------------- +
   Master mix for one sample:
   4 µl NEBuffer3.1 10x buffer (keep on ice)
   1 µl NEB RE 1 ex AseI (Or any other RE of choice)
   1 µl NEB RE 2 ex NsiI (Or any other RE of choice)
   4 µl MilliQ
   ----------------------------
   40 µl total

2. Add 10 µl of the master mix to each sample.
3. Spin and Vortex the samples
4. Perform the digestion at 37°C for 17 hours (using the thermal heating-block).
   Note: The incubation time is longer than what is recommended by the RE manufacturers. However in our experience longer incubation gave better DNA yield for the preparation of epiGBS libraries.
5. Once digestion is completed, samples can be stored at 4°C for several days.

#### 3.2 Annealing Adapters

Note: This step only needs to be done once, when using the adapters for the first time.

1. Mix

   50 µl BA adapter I (100 microM) or CO adapter I (100 µM)
   50 µl BA adapter II (100 microM) or CO adapter II (100 µM)
   ------------------------------------
   100 µl total (50 pmol/µl) = 50 µM

2. Perform the annealing:

   5 min 95 °C
   each minute -1 °C x 70 times (end temperature = 25 °C)
   5 min 4 °C

3. Use 6 µl of the annealed adapter concentrate for the first dilutions and store the rest at -20 °C.
4. Add 6 µl concentrated annealed stock to 1 ml  5 mM Tris/HCl, pH 8.5. The resulting concentration varies from ~3-10 ng / µl dsDNA.
5. Measure the concentration using fluorescence and freeze the leftover diluted stock at -20 °C.
6. Further dilute the diluted stock using x µL (calculate per sample) in 500 µl 5 mM Tris/HCl, pH 8.5 to 600 pg / µl.
7. Make a working stock by aliquoting the final dilution in 100 µl portions.
8. Freeze all aliquots except the one in use, which can be stored for 2 months at 4 °C.

Avoid freezing-thawing cycles as much as possible!

#### 3.3 Adapter Ligation

- Beforehand: Remove the T4 DNA ligase buffer and adapters from the freezer and allow them to thaw on ice.

1. Add the following in 0.5 ml PCR reaction tubes in this order: The digested DNA, then the corresponding barcodes, then 20 µl of a ligation master mix consisting of T4 DNA ligase buffer, T4 DNA ligase and MiliQ water.

| | | |
|---|---|---|
| 40 | µl | Digested DNA |
| -------------------- + |  |  |
| Master mix for one sample: | | |
| 6 | µl | T4 DNA Ligase buffer 10x |
| 0.5 | µl | T4 DNA ligase |
| 4 | µl | Barcoded BA adapter (600 pg / µl) |
| 4 | µl | Barcoded CO adapter (600 pg / µl) |
| 5.5 | µl | MilliQ |
| ----------------------------- |  |  |
| 60 | µl | total |

2. Perform the ligation, at 22 °C block temperature and 30 °C lid temperature in a PCR machine for 17 hours

3. Samples can be stored for several days at 4°C

#### 3.4 Multiplexing

Only samples that A) were digested by the same RE combination and B) originated from the same species should be multiplexed.

1. Spin, individually, all digested and adapter-ligated DNA samples.
2. Combine all of the DNA samples into a single master DNA pool.
3. Vortex the multiplexed master DNA pool.
4. It is advised to store 1/3 of the total volume of the multiplexed master DNA pool at -20°C as backup.
5. Divide the remaining 2/3 of the multiplexed master DNA pool into several 2 ml reaction tubes. These tubes will be used as technical replicates. The number of technical replicates vary depending on the total number of samples multiplexed together. It is advised that each technical replicate contains symbolically 12 samples, e.g. if 96 samples are multiplexed, 96/12 = 8 technical replicates. If 72 samples are multiplexed, 72/12 = 6 technical replicates. This adds up to, on average, 0.5 ml of multiplexed master DNA pool per technical replicate. These technical replicates will control for any biases that are created during the bisulfite (BS) conversion. It also ensures that the columns used during the cleaning and BS steps are not overloaded.
6. Quantify the concentration of the multiplexed master DNA pool on a Qubit fluorometer.

#### 3.5 Concentrating and Cleaning

- This step was adapted from the NucleoSpin Gel & PCR Cleanup Kit protocol.
- Beforehand: Pre-heat the NE elution buffer at 55°C in the microplate shaker at maximum rpm.

In each technical replicate:

1. Add 10 µl of 3M NaAc pH 5.3 and 1.2 ml NTI buffer. The NaAc is added to adjust the pH of the binding buffer. The buffer should turn from slightly orange to yellow.
2. Vortex and centrifuge at 11 000 g for 30 seconds.
3. Add 600 µl of the mix to the provided NucleoSpin Clean-up column, centrifuge at 11000 g for 1 min and remove the flow-through.
4. Repeat the previous step by adding another 600 µl of the mix and then repeat a third time by adding the remaining mix.

5. Wash the column with 600 µl NT3 wash buffer, centrifuge at 11000 g for 1 min and remove the flow-through.
6. Repeat the previous step.
7. Centrifuge at 11 000 g for 30 seconds and place the column into a new 1.5 ml reaction tube.
8. Incubate the column in the reaction tube (with the lid open) in the microplate shaker at 55°C for 5 min.
9. Elute the DNA with 40 µl of NE-elution buffer into a new 2.0 ml Lobind reaction tube. Wait 1 min at RT and then centrifuge at 11 000 g for 1 min.
10. Repeat the previous step with the 40 µl of eluted DNA.
11. Determine the DNA concentration with a Qubit Fluorometer.

#### 3.6 0.8X SPRI size-selection

- Beforehand:
  - Remove SPRIselect beads from the fridge and store them at RT. Before usage, thoroughly shake the SPRIselect bottles to resuspend SPRI beads.
  - Prepare fresh 70 % ethanol. For 8 technical replicates, you will need around 10 ml (7.3 ml 96 % ethanol with 2.7 ml MiliQ water).
  - Calculate the amount of SPRIselect beads solution that you will need to add to each technical replicate:
    - Determine the exact total volume of eluted DNA in the reaction tubes (this should be around 40 µl)
    - Volume of sample X SPRIselect bead ratio = Volume of SPRIselect beads solution, e.g. 40 µl X 0.8xratio = 32 µl of SPRIselect.

1. Add the calculated volume of SPRIselect to each corresponding technical replicate.
2. Mix the total volume by pipetting the mix 10 times up and down and incubate at RT for 1 minute.
3. Place technical replicates, one at a time, on the magnetic rack.
   Note: We recommend working with a maximum of 3 technical replicates at a time on the magnetic rack to reduce the amount of time that the SPRIselect beads are manipulated in each technical replicate.
4. Wait for the beads to fully separate, forming a dark pellet at the bottom of the reaction tube.
5. Remove the transparent supernatant, ensuring not to disrupt the pellet.

Note: It is advised to leave the technical replicates on the magnetic rack while removing the transparent flow-through.

6. Quickly add 500 µl of freshly prepared 70 % ethanol onto the SPRIselect beads and resuspend the beads by pipetting up and down.

   Note: It is important not to wait too long before adding the 70 % Ethanol to the SPRIselect beads as the SPRIselect bead pellet will dry, which makes the resuspension harder.
7. Repeat step 4-5
8. Quickly add 600 µl of 70 % ethanol to the beads and resuspend the SPRIselect beads by pipetting up and down.
9. Wait for SPRIselect beads to form a pellet and remove the transparent supernatant.
10. Allow for the SPRIselect beads to dry three to five minutes at RT.
11. When dry, remove the technical replicates from the magnetic rack and add 23 µl of NE Elution Buffer onto the SPRIselect beads. Incubate for 3 minutes at RT and then resuspend beads by pipetting up and down.
12. Place the technical replicates back onto the magnetic rack, wait for the beads to form a pellet and transfer the transparent supernatant into a new 2 ml reaction tube.
13. Repeat the cleaning step until all technical replicates are done.
14. Determine the DNA concentration on Qubit Fluorometer.

#### 3.7 Nick Repair

- Beforehand: Defrost the NEB2 Buffer and 5-Methylcytosine dNTPs on ice

1. In 0.5 ml PCR tubes add 19.25 µl cleaned DNA and then 5.75 µl of a master mix consisting of NEB2 buffer, 5-Methylcytosine dNTPs and DNA Polymerase I:

   19.25 µl DNA
   ---------------------- +
   Master mix for one technical replicate:
   2.5 µl NEB2 buffer
   2.5 µl 5-Methylcytosine dNTP's
   0.75 µl DNA Polymerase I
   -----------------------------
   25 µl Total

2. Perform the Nick repair in a PCR machine for 1 hour, with a block temperature of 15 °C and a lid temperature of 30 °C.

#### 3.8 Test PCR (Optional)

The aim of this step is to test that the primers are correctly annealing, thus ensuring maximal performance during the epiGBS PCR in step 3.10 . If using a new species, this step can also be used to determine the optimal amount of PCR cycles needed to amplify the DNA (See Boquete *et al.,* 2020 for more information).

- Beforehand:
    - Thaw the KAPA HiFi Hot Start* Readymix PCR kit and Primers on ice.
    - If necessary, dilute the stock Primers (100 µM) to a working solution with a final concentration of 10 µM.

1. Add 1 µl DNA to a 0.5 ml PCR tube and add 9 µl of a master mix consisting of Kapa Hifi HS+ReadMix PCR, Primer A, Primer B and MiliQ water:

 1 µl DNA
---------------------- +
Master mix for one technical replicate:
 5 µl KAPA HIFI HS + ReadyMix PCR Kit
0.3 µl Primer A (10 µM)
0.3 µl Primer B (10 µM)
3.4 µl MilliQ
-----------------------------
10 µl Total

2. Execute the PCR using the following program:

95 °C 3 minutes
-------------------------------------------------------------------------
98 °C 10 seconds
65 °C 15 seconds 14 cycles
72 °C 15 seconds
-------------------------------------------------------------------------
72 °C 5 minutes

3. Determine DNA concentration on a Qubit fluorometer.

#### 3.9 Bisulfite Conversion (BS)

Note: It is recommended to follow the protocol from the EZ DNA Methylation-Lightning Kit from Zymo (Catalogue Number: D5030).

1. To 20 µl of nick-repaired DNA of each technical replicate, add 130 µl Lightning Conversion Reagent( this should correspond to 6.5 µl lightning conversion reagent/µl of DNA) in a 0.5 ml PCR tube.
   Note: if the volume of DNA is less than 20 µl, add MiliQ water.
2. Vortex and centrifuge at 11000 g for 30 seconds.
3. Place the PCR tube in a PCR machine and run the following program:

   98 °C 8 minutes
   54 °C 60 minutes

   Note: DNA can then be stored at 4°C for up to 20 hours.

4. Apply 600 µl M-Binding Buffer to a Zymo-Spin TM IC Column and place the column into the provided collection tube.
5. Apply the DNA to the Zymo-Spin TM IC Column containing the M-Binding Buffer. Close the cap and mix by inverting the column 5 times.
6. Centrifuge at 11000 g for 30 seconds. Discard the flow-through.
7. Apply 100 µl of M-Wash Buffer to the column. Centrifuge at 11000 g for 30 seconds and discard the flow-through.
8. Apply 200 µl of M-Wash Buffer to the column. Centrifuge at 11000 g for 30 seconds.
9. Repeat this last wash step before discarding the flow-through.
10. Place the column into a 1.5 ml microcentrifuge tube and add 12 µl M-Elution Buffer directly to the column matrix. Centrifuge at 11000 g for 30 seconds to elute the DNA.

    Note: At this stage, the DNA can be stored for a longer period of time. It is recommended that you store it at -20 °C for short-term storage or at -80 °C for long-term storage.

#### 3.10 Library amplification PCR

- Beforehand:
  - Thaw the KAPA HiFI HS Uracil+ ReadyMix PCR kit and Primers on ice.
  - Depending on the DNA concentration of each technical replicate, it is recommended to use two to five PCR reactions per technical replicate to increase the amplification efficiency.

1.
   a. For 1X reaction, add 8 µl of master mix consisting of KAPA HiFi HS Uracil+ ReadMix PCR, Primer A, Primer B and MiliQ water to 2 µl of BS converted DNA in a 0.5 ml PCR tube:

      2 µl DNA
      --------------------- +
      Master mix for one technical replicate:
      5 µl KAPA HIFI Uracil+ hotstart ready mix
      0.3 µl Primer A (10mM)

0.3 µl Primer B (10mM)
2.4 µl MiliQ
----------------------------
10 µl total

b. For 5X reaction, add 40 µl of a master mix consisting of KAPA HiFi HS Uracil+ ReadMix PCR, Primer A, Primer B and MiliQ water to 10 µl of BS converted DNA in a 0.5 ml PCR tube:

10 µl DNA
--------------------- +
Master mix for each technical replicate:
25 µl KAPA HIFI Uracil + hotstart ready mix
1.5 µl Primer A (10 mM)
1.5 µl Primer B (10 mM)
12 µl MilliQ (can also be added during extra BS elution step)
----------------------------
50 µl total

2. Execute the PCR using the following PCR program:

95 °C 3 minutes
----------------------------------
98 °C 10 seconds
65 °C 15 seconds 14-16 cycles**
72 °C 15 seconds
----------------------------------
72 °C 5 minutes

** The number of cycles will depend on the results of the test PCR (See point 3.8 and Boquete *et al.*, 2020). Generally, 14 cycles is enough to amplify the DNA.

3. Determine the DNA concentration on a Qubit fluorometer.

#### 3.11 Library Cleaning

As for step 3.5, this step was adapted from the Nucleospin Gel & PCR Cleanup Kit protocol.

- Beforehand:
  - Pre-heat the NE elution buffer at 55°C in the microplate shaker at maximum rpm.

- Combine 2 technical replicates into a 1.5 ml new reaction tube. In other words, if you work with 8 technical replicates, 4 technical replicate pools will remain. The expected volume per reaction tube is 100 µl.

To each technical replicate pool:

1. Add 200 µl NTI buffer and 1 µl 3M NaAc pH 5.3.
2. Vortex and centrifuge at 11000 g for 30 seconds.
3. Add 700 µl of the mix to the provided NucleoSpin Clean-up column, centrifuge at 11000 g for 1 min, remove the flow-through.
4. Repeat step 3 three times.
5. Wash the column with 600 µl NT3 wash buffer, centrifuge at 11000 g for 30 seconds.
6. Repeat step 5 step twice.
7. Transfer the column into a new 1.5 ml reaction tube and centrifuge at 11000 g for 30 seconds.
8. Incubate the column in the reaction tubes (with the lid open) in the microplate shaker at 55°C for 5 min.
9. Elute the DNA with 50 µl of EB-elution buffer in a new 2.0 ml Lobind reaction tube. Wait 1 min at RT and then centrifuge at 11000 g for 1 min.
10. Repeat the previous step using the 50 µl eluted DNA.
11. Determine the DNA concentration with a Qubit fluorometer.

#### 3.12 0.8X SPRI size-selection

- Beforehand:
  - Remove SPRIselect beads from the fridge and store them at RT. Before usage, thoroughly shake the SPRIselect bottles to resuspend SPRI beads.
  - Prepare fresh 70 % ethanol.
  - Calculate the amount of SPRIselect beads solution that you will need to add to each technical replicate:
    - Determine the exact total volume of eluted DNA in the reaction tubes (this should be around 40 µl)
    - Volume of sample X SPRIselect bead ratio = Volume of SPRIselect beads solution ex 40 µl X 0.8xratio = 32 µl of SPRIselect.

1. Add the calculated volume of SPRIselect to each corresponding technical replicate pool.

2. Mix the total volume by pipetting the mix 10 times up and down and incubate at RT for 1 minute.
3. Place technical replicate pools, one at a time, on the magnetic rack.
   Note: We recommend working with a maximum of 3 technical replicate pools at a time on the magnetic rack reducing the amount of time the SPRIselect beads are being manipulated in each technical replicate pool.
4. Wait for the beads to fully separate, forming a dark pellet at the bottom of the reaction tube.
5. Remove the transparent supernatant, ensuring not to disrupt the pellet.
   Note: It is advised to leave the technical tubes on the magnetic rack while removing the transparent supernatant.
6. Quickly add 500 µl of freshly prepared 70 % ethanol onto the SPRIselect beads and resuspend the beads by pipetting up and down.
   Note: It is important not to wait too long before adding the 70 % Ethanol to the SPRIselect beads as the SPRIselect bead pellet will dry rendering the resuspension harder.
7. Repeat step 4-5
8. Quickly add 600 µl of 70 % ethanol to the beads and resuspend the SPRIselect beads by pipetting up and down.
9. Wait for SPRIselect beads to form a pellet and remove the transparent supernatant.
10. Allow for the SPRIselect beads to dry three to five minutes at RT.
11. When dry, remove the technical replicate pools from the magnetic rack and add 23 µl of NE Elution Buffer onto the SPRIselect beads. Incubate for 3 minutes at RT and then resuspend beads by pipetting up and down.
12. Place the technical replicate pools back onto the magnetic rack, wait for the beads to form a pellet and transfer the transparent supernatant into a new 2 ml reaction tube.
13. Repeat the cleaning step until all technical replicate pools are done.
14. Determine the DNA concentration on Qubit Fluorometer.

#### 3.13 Library Quality Check

- DNA concentration of final library using qPCR:
  1. Combine all technical replicate pools into one reaction tube.

2. We recommend following the KAPA Library Quantification for NGS (Roche Sequencing, Catalogue Number: KK4844).

   Note: Usually, 3 ng/µl final epiGBS library DNA is sufficient for sequencing.

- Library fragment Analyzing using a Bio-fragment Analyser:
  1. We recommend following the High Sensitivity NGS Fragment Analysis (1-6000bp) kit (Advance Analytical, Catalogue Number: DNF-473-5000)

     Note: The fragment size expected is around 400 bp.

#### 3.14 Sequencing

- The epiGBS protocol was optimized to run on a single lane of a HiSeq X 150 bp PE (Illumina) sequencing machine.
- Spike 12 % phiX.

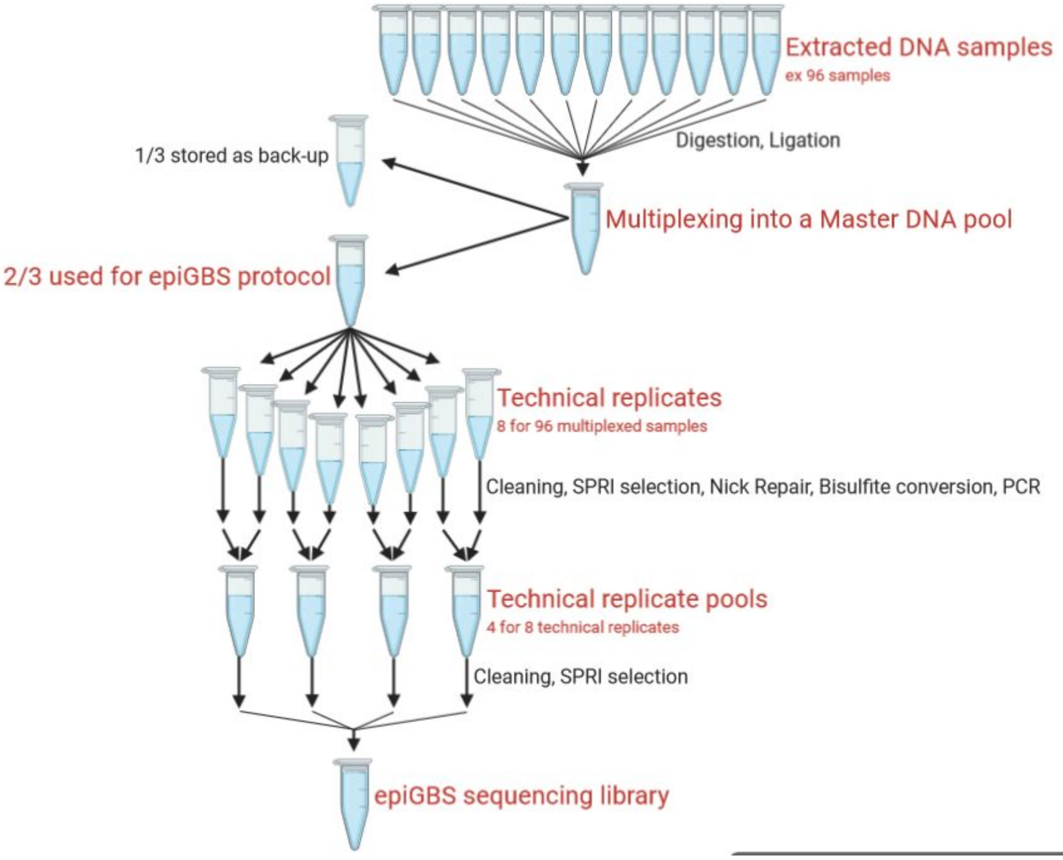

Figure 1: Representation of the various demultiplexing and pooling steps occurring in the epiGBS2 laboratory protocol. It is suggested to start working with a multiple of 12 samples (e.x 96, 84, 72, 60… samples), in order to keep a correct ratio of samples per pool when demultiplexing and/or pooling.
