## Supplementary material for "epiGBS2: an improved protocol and automated snakemake workflow for highly multiplexed reduced representation bisulfite sequencing": report with example data: report-example.html

epiGBS report


### epiGBS report

###### author: morganea

###### date: 2020-04-06

This report contains all important information about your epiGBS analysis. The ouput of this analysis is stored in `/home/morganea/epiGBS2/output`. This report will give you a quick overview of the quality of your results. For more details please always refer to the original (log)files:

| File | path |
| --- | --- |
| bedfile | `/home/morganea/epiGBS2/output/mapping/methylation.bed` |
| snpfile | `/home/morganea/epiGBS2/output/mapping/snp.vcf.gz` |
| multiQC-report | `/home/morganea/epiGBS2/output/multiQC_report.html` |
| demultiplexing-log | `/home/morganea/epiGBS2/output/output_demultiplex/demultiplex.log` |
| stacks-log | `/home/morganea/epiGBS2/output/output_demultiplex/clone-stacks/process_radtags.clone.log` |
| mapping-log | `/home/morganea/epiGBS2/output/mapping/mapping_variantcalling.log` |
| denovo-log | `/home/morganea/epiGBS2/output/output_denovo/make_reference.log` |

### Tabset

#### Clones Removal

The PCR clones are removed from the raw sequencing reads using a random nucleotide sequence (“wobble”) in the adapters.

Before removal, your raw reads contained 12.43% clone reads.

The histogram shows the distribution of PCR clone numbers. Ideally, most reads will occur once “1”.

Fig. 1: Distribution of clone numbers

#### Demultiplexing

You performed demultiplexing with the process\_reads program from stacks. The following stats table will give you information about how many reads were retained after barcode- and RAD-tag (RE cut site) check. For the RAD-tag check 1 mismatch is allowed to correct for mismatches due to C/T conversion in the RAD-tag. Check your barcode or your enzyme information if the amount of retained reads is suprisingly low.

```
## 831669476 total sequences
## 178481716 ambiguous barcode drops (21.5%)
##         0 low quality read drops (0.0%)
##  27417780 ambiguous RAD-Tag drops (3.3%)
## 625769980 retained reads (75.2%)
```

In a well designed study, read numbers per sample should be similar. You can check in the following histogram, if this is true for your current analysis. The vertical, dotted line indicates the mean. Please also check the multiQC stats for a full overview about the read quality.

Fig.2: Histogram of the number of demultiplexed reads per read file

The (non-)conversion rate is a measurement for the efficiency of your BS-treatment. Sequences with a control nucleotide “C” in the forward (R1) read and a “T” in the reverse (R2) read are normally called “Watson”, vice versa “Crick”. Sequences with a “C” in both R1 and R2 reads are considered as non-converted.

```
## [1] "You have got 309031345 Watson reads, 316738635 Crick reads and 11536362 non-converted reads. Your non-conversion rate is 1.81 %."
```

Also check the multiQC stats for a full overview about the read quality.

#### De novo reference

During demultiplexing all reads were split into samples and annotated as Watson or Crick strands. This information is used during creation of the de novo reference clusters.

During the first step Watson and Crick forward and reverse reads are or assembled (merged), if they overlap, or joined by adding poly-N, respectively. A low assembly % might indicate a low read quality (on the end of the reads) or an insert size longer than twice your read size.

You assembled reads, when they met the following parameters:

```
## Minimum overlap....................: 10
## p-value............................: 0.001000
```

The percentage of assembled and un-assembled (joined reads) was:

Crick first, followed by Watson.

```
## Assembled reads ...................: 135,057,335 / 153,494,118 (87.989%)
## Discarded reads ...................: 0 / 153,494,118 (0.000%)
## Not assembled reads ...............: 18,436,783 / 153,494,118 (12.011%)
##
## Assembled reads ...................: 129,270,398 / 146,541,950 (88.214%)
## Discarded reads ...................: 0 / 146,541,950 (0.000%)
## Not assembled reads ...............: 17,271,552 / 146,541,950 (11.786%)
```

Assembly and joining of forward and reverse reads is followed by three different clustering steps. In the first step the Watson and Crick reads are deduplicated. Afterwards the deduplicated reads are binarized and clustering is performed to pair binary Watson and Crick reads. The original sequence is restored, followed by clustering based on identity. The identity % can be set in the snakemake config file.

It follows the stats for the first clustering step (deduplication).

Watson-joined:

```
## 5008750080 nt in 17271552 seqs, min 290, max 290, avg 290
## 6977812 unique sequences, avg cluster 2.5, median 1, max 169130
```

Watson-assembled:

```
## 16403179692 nt in 127725950 seqs, min 32, max 271, avg 128
## minseqlength 32: 1544448 sequences discarded.
## 11318639 unique sequences, avg cluster 11.3, median 1, max 3702794
```

Crick-joined:

```
## 5346667070 nt in 18436783 seqs, min 290, max 290, avg 290
## 9187311 unique sequences, avg cluster 2.0, median 1, max 169611
```

Crick-assembled:

```
## 17250341921 nt in 133406532 seqs, min 32, max 271, avg 129
## minseqlength 32: 1650803 sequences discarded.
## 13449210 unique sequences, avg cluster 9.9, median 1, max 4311502
```

It follows the stats of the second clustering step (uc):

```
## 9282533885 nt in 40932971 seqs, min 32, max 290, avg 227
## 31918267 unique sequences, avg cluster 1.3, median 1, max 4645
```

It follows the stats of the third clustering step (identity). Usually you will loose the majority of clusters from step 2, because most clusters do not consist of Watson and Crick deduplicated reads but only Watson or only Crick. Those clusters will be excluded:

```
## 308657197 nt in 1490797 seqs, min 32, max 290, avg 207
## Clusters: 146745 Size min 1, max 23592, avg 10.2
## Singletons: 52190, 3.5% of seqs, 35.6% of clusters
```

#### Mapping

To determine variants (SNPs or methylation), sequencing reads are mapped against the de-novo reference. Mapping is performed with STAR. A low mapping percentage might indicate low quality of the sequencing reads (e.g. 3’ adapter sequences), a non-suitable reference or problems with the de novo sequence creation. The higher the % Uniquely mapped reads, the higher the quality of your alignment is. A high % of reads mapped to multiple or too many loci indicates that you have or very low complexity reads or typically many repetitive regions in your reference (% identity in de novo reference creation too high). % of reads unmapped: too many mismatches might indicate that % identity in the de novo sequence creation might be too low. % of reads unmapped: too short means that reads could not properly aligned to the reference (but does (confusingly) NOT indicate that reads are actually too short)

It follows the mapping stats:

Watson-joined:

```
##                         Uniquely mapped reads % |    79.20%
##              % of reads mapped to multiple loci |    0.00%
##              % of reads mapped to too many loci |    1.27%
##        % of reads unmapped: too many mismatches |    8.96%
##                  % of reads unmapped: too short |    7.66%
##                      % of reads unmapped: other |    2.91%
```

Watson-assembled:

```
##                         Uniquely mapped reads % |    49.08%
##              % of reads mapped to multiple loci |    0.00%
##              % of reads mapped to too many loci |    32.57%
##        % of reads unmapped: too many mismatches |    1.04%
##                  % of reads unmapped: too short |    3.41%
##                      % of reads unmapped: other |    13.90%
```

Crick-joined:

```
##                         Uniquely mapped reads % |    76.02%
##              % of reads mapped to multiple loci |    0.00%
##              % of reads mapped to too many loci |    0.94%
##        % of reads unmapped: too many mismatches |    12.12%
##                  % of reads unmapped: too short |    8.11%
##                      % of reads unmapped: other |    2.80%
```

Crick-assembled:

```
##                         Uniquely mapped reads % |    48.57%
##              % of reads mapped to multiple loci |    0.00%
##              % of reads mapped to too many loci |    32.41%
##        % of reads unmapped: too many mismatches |    1.25%
##                  % of reads unmapped: too short |    3.57%
##                      % of reads unmapped: other |    14.19%
```

#### SNP calling

The epiGBS pipeline performs SNP calling. The SNP depth reflects the reliability of a called SNP and depends e.g. from the amount of reads and the mappibility of the reads.

In total, you have got 2021404 SNPs.

The figure shows the average SNP depth over all samples.

Fig.3: Histogram of average SNP depth over all samples

#### Methylation calling

This might be the part, you are interested in the most. You probably will continue your statistical analysis with the methylation.bed file. Here you will find some simple summarized stats about the methylation calls. Please realize, that all following plots are based on the first 100000 methylation sites because of memory issues. You originally obtained 10417837 methylation sites.

The following histogram shows the distribution of the number of samples, in which methylated positions were called.

Fig.5: Distribution of the number of samples, that have a specific methylation site called

The next bar diagram shows, how many sites you obtained in each context CG, CHG, CHH.

Fig. 6: Number of methylated sites in each context

The following diagram shows the distribution of the methylation site depth per samples. The x-axis is limited to 250. The vertical line indicated the median.

```
## (0, 250)
```

Fig.7: Distibution of the average depth per methylation site and sample

The following diagram shows the distribution of methylation ratio (“methylated”/“total”) in different contexts (CG, CHG, CHH)

Fig.8: Distribution of methylation ratio per context
